# Mitochondrial-Derived Compartments Remove Surplus Proteins from the Outer Mitochondrial Membrane

**DOI:** 10.1101/2023.07.07.548175

**Authors:** Zachary N. Wilson, Sai Sangeetha Balasubramaniam, Mitchell Wopat, Adam L. Hughes

**Affiliations:** Department of Biochemistry, University of Utah School of Medicine, _Salt Lake City_, UT 84112, USA; Lead contact

## Abstract

The outer mitochondrial membrane (OMM) creates a boundary that imports most of the mitochondrial proteome while removing extraneous or damaged proteins. How the OMM senses aberrant proteins and remodels to maintain OMM integrity remains unresolved. Previously, we identified a piecemeal autophagic mechanism called the <u>m</u>itochondrial-<u>d</u>erived <u>c</u>ompartment (MDC) that removes a subset of the mitochondrial proteome. Here we show that MDCs specifically sequester proteins localized only at the OMM, providing an explanation for how select mitochondrial proteins are removed by MDCs. Remarkably, selective sorting into MDCs also occurs within the OMM, as subunits of the translocase of the outer membrane (TOM) complex are excluded from MDCs unless assembly of the TOM complex is impaired. Considering that overloading the OMM with mitochondrial membrane proteins or mistargeted tail-anchored membrane proteins induces MDCs to form and sequester these proteins, we propose that one functional role of MDCs is to create an OMM-enriched trap that segregates and sequesters excess proteins from the mitochondrial surface.

**SUMMARY:** Wilson and colleagues observe that mitochondrial-derived compartments (MDCs) selectively incorporate proteins from only the outer mitochondrial membrane (OMM), and robustly sequester both excess and mistargeted proteins into this OMM-enriched domain, suggesting MDCs act to remove surplus hydrophobic cargo from mitochondria.

## INTRODUCTION

Mitochondria are double-membrane bound organelles that perform critical roles in cell physiology. While the inner mitochondrial membrane (IMM) performs essential roles in cell metabolism, the outer mitochondrial membrane (OMM) creates a key interface that supports mitochondrial biogenesis and establishes mitochondrial connections throughout the cell (Pfanner et al., 2019; Harper et al., 2020). Because 99% of the mitochondrial proteome is coded within nuclear genes, the OMM contains several protein import complexes and targeting receptors that collaborate with cytosolic chaperones to capture mitochondrial precursor proteins and direct them into mitochondria (Wiedemann and Pfanner, 2017). Failing to capture mitochondrial precursor proteins can cause dramatic consequences for the cell, leading mitochondrial proteins to aberrantly target to other organelles (Vitali et al., 2018; Xiao et al., 2021; Shakya et al., 2022) or to accumulate and aggregate, leading to mitoprotein-induced stress responses (Wang and Chen 2015, Wrobel et al., 2015; Boos et al., 2019). Conversely, the impaired import of mitochondrial precursor proteins induces the action of several mitochondrial quality control pathways that collaborate with the cytosolic ubiquitin-proteasome system to safeguard the integrity of the OMM. These pathways act by recruiting AAA-ATPases that use the energy from ATP hydrolysis to remove stalled, misfolded, or damaged proteins from the OMM and direct these proteins to cytosolic proteasomes for degradation (Heo et al., 2010; Weidberg and Amon, 2018; Mårtensson et al., 2019, Metzger et al., 2020).

The fidelity of mitochondrial protein targeting and import is further challenged by similarities in protein targeting mechanisms that occur at other organelles (Hegde and Keenan, 2021). The ER is a major biosynthetic organelle responsible for importing and folding ∼30% of the cellular proteome, receiving and importing precursor proteins through both co-translational and post-translational mechanisms (Juszkiewicz and Hegde, 2018). Similar to mitochondrial protein targeting, the loss of correct protein targeting to the ER can lead to some proteins, particularly membrane proteins, to mistarget to the mitochondrial OMM (Schuldiner et al., 2008; Chen et al., 2014; Costa et al., 2018). For example, in the absence of the guided entry of tail-anchored proteins (GET) pathway, tail-anchored (TA) membrane proteins mislocalize, generating cytosolic protein aggregates or mistarget to the OMM (Schuldiner et al., 2008; Li et al., 2019). To protect against these mistargeted TA proteins, mitochondria contain a membrane-embedded AAA-ATPase, ATAD1/Msp1, that directly removes mistargeted TA proteins and delivers these proteins for proteasomal degradation or retargeting (Chen et al., 2014; Wohlever et al., 2017; Li et al., 2019; Matsumoto et al., 2019). While both mitochondria and the ER contain mechanisms to remove aberrant or damaged proteins, the accumulation of aberrant proteins on the surface of these organelles can also induce dramatic membrane remodeling to preserve organelle homeostasis.

The accumulation of membrane proteins within the ER induces a form of ER microautophagy, whereby the ER membrane proliferates, creating stacked membrane sheets and spherical ER whorls that robustly sequester certain ER membrane proteins and delivers them to vacuoles/lysosomes for degradation (Wright et al., 1987; Schuck et al., 2009; Schäfer et al., 2020). In mammalian cells, mitochondrial failure and impaired mitochondrial import can lead to the accumulation of PINK1 on the mitochondrial surface (Lazarou et al., 2012; Okatsu et al., 2013), initiating a signaling cascade that leads to the wholesale removal of entire mitochondria (Pickles et al., 2018). Conversely, rather than the removal of entire organelles, piecemeal degradative mechanisms can facilitate the removal of select portions of mitochondria in a manner that is kinetically faster and less energy intensive. In response to some stress conditions, particularly mild oxidative damage, mitochondria form small vesicles (60-150nm diameter) that incorporate damaged proteins and delivers them to lysosomes for degradation (Soubannier et al., 2012). These mitochondrial-derived vesicles (MDVs) have been shown to compensate for the loss of mitophagy and may be an early response pathway that attempts to protect the mitochondrial network prior to the onset of mitophagy (McClelland et al., 2014; Towers et al., 2021). Interestingly, mitochondria can also be induced to shed their outer membrane in response to infection-induced stress elicited by the targeting of pathogen proteins to the OMM (Li et al., 2022). Contrary to MDVs, these infection-induced OMM proliferations are large (several microns in diameter) and appear to contain internal membrane invaginations, somewhat reminiscent to ER whorls (Li et al., 2022).

Previously, in the budding yeast, *Saccharomyces cerevisiae*, we identified a mitochondrial quality control pathway that also involves the remodeling of mitochondrial membranes to form a mitochondrial-derived compartment (MDC). In old-aged yeast cells and in response to acute stressors that alter cellular amino acid and lipid homeostasis, mitochondria form large (near micron-sized), spherical compartments that robustly sequester only a minor portion of the mitochondrial proteome (Hughes et al., 2016; Schuler et al., 2021; Xiao et al., 2023). MDCs are generated from dynamic mitochondrial membrane extensions that repeatedly elongate, coalesce, and invaginate to generate distinct mitochondrial domains that robustly sequester select mitochondrial proteins (English et al., 2020; Wilson et al., 2023 *preprint*). Subsequently, MDCs are removed from mitochondria and delivered to yeast vacuoles for degradation, suggesting that MDCs act as a piecemeal autophagic mechanism that is induced to remodel or ensnare a portion of the mitochondrial proteome (Hughes et al., 2016). The primary cargoes identified within MDCs all include mitochondrial membrane proteins or membrane-associated proteins, including the OMM surface receptor Tom70 and members of the mitochondrial metabolite carrier superfamily (Hughes et al., 2016). While mitochondrial metabolite carriers are incorporated into MDCs, the majority of other IMM proteins, as well as, aqueous proteins from the mitochondrial matrix or the intermembrane space were all excluded from MDCs (Hughes et al., 2016). How protein cargoes are specifically sorted into MDCs, where these cargo proteins are sorted from, and the purpose for this segregation remains unclear.

Through an examination of protein selectivity into MDCs we show here that MDCs specifically sequester proteins that are only localized at the OMM, providing an explanation for how MDCs incorporate a minor subset of the mitochondrial proteome. Intriguingly, cargo sequestration into MDCs is also selective within the OMM, as subunits of the TOM complex are precluded from entering MDCs unless TOM complex assembly is impaired. Prompted by these results and the observation that MDCs sequester several layers of OMM (Wilson et al., 2023 *preprint*), we determined that MDCs form in response to the aberrant accumulation of OMM proteins, including proteins mistargeted to the OMM. By following the mistargeting of TA proteins to the OMM, we show that MDCs can selectively sequester mistargeted TA proteins and thus may act in concordance with other mitochondrial quality control pathways to preserve OMM integrity.

## RESULTS

### N-terminally tagged mitochondrial metabolite carriers are excluded from MDCs

Previously we used the yeast GFP collection to identify the cohort of mitochondrial proteins that are sequestered into MDCs. Out of 304 detectable mitochondrial proteins that are C-terminally fused to GFP, only 26 proteins were identified to enter MDCs (Hughes et al., 2016). All of these proteins were integral membrane proteins or membrane-associated proteins from both the outer and inner mitochondrial membranes. A distinct class of proteins that preferentially incorporated into MDCs were mitochondrial metabolite carriers (Hughes et al., 2016). Because we observed that MDCs form in response to certain metabolic stressors (Schuler et al., 2021), we hypothesized that specific intrinsic features of mitochondrial metabolite carriers are important for their selective sorting into MDCs. To analyze how mitochondrial metabolite carriers are incorporated into MDCs, we used the yeast oxaloacetate carrier, Oac1, as a prototype for MDC cargo sequestration.

Oac1 represented an ideal cargo for examining protein selection into MDCs, as Oac1-GFP readily incorporates into MDCs (Hughes et al., 2016; Schuler et al., 2021) and Oac1 participates in leucine biosynthesis providing a means to assess Oac1 function (Fig. 1 A; Kohlhaw 2003; Marobbio et al., 2008). A double deletion of the *LEU4* and *OAC1* genes created a yeast strain (*leu4*Δ*oac1*Δ) that is auxotrophic for leucine and growth can be rescued by ectopically expressing Oac1 from a low or high-copy plasmid (Fig. 1 B; Marobbio et al., 2008). Using this analysis, we determined that Oac1 C-terminally fused to GFP produces a non-functional protein as it cannot rescue the growth of the *leu4*Δ*oac1*Δ strain when grown on medium lacking leucine (Fig. 1 B). Alternatively, placing GFP at the N-terminus of Oac1 created a functional protein as it rescued growth of the *leu4*Δ*oac1*Δ strain when expressed from a low or high-copy plasmid (Fig. 1 B). Considering both constructs localize to mitochondria, we compared these two differentially GFP-tagged versions of Oac1 for incorporation into MDCs, which can be induced to form by treating yeast with rapamycin (Schuler et al., 2021). Intriguingly, while Oac1-GFP readily incorporates into MDCs (the domains for which are marked by the strong enrichment of Tom70-mCherry (Fig. 1 C, white arrows)), GFP-Oac1 is robustly excluded from MDCs (Fig. 1, C and D). The minor percentage (<20%) of GFP-Oac1 we observed at MDCs can be attributed, in part, to our scoring technique, where we first assigned MDCs based on Tom70-mCherry enrichment (which occasionally produces false MDC identifications) and where some MDCs contained a faint but detectable level of GFP-Oac1.

**Figure 1.**
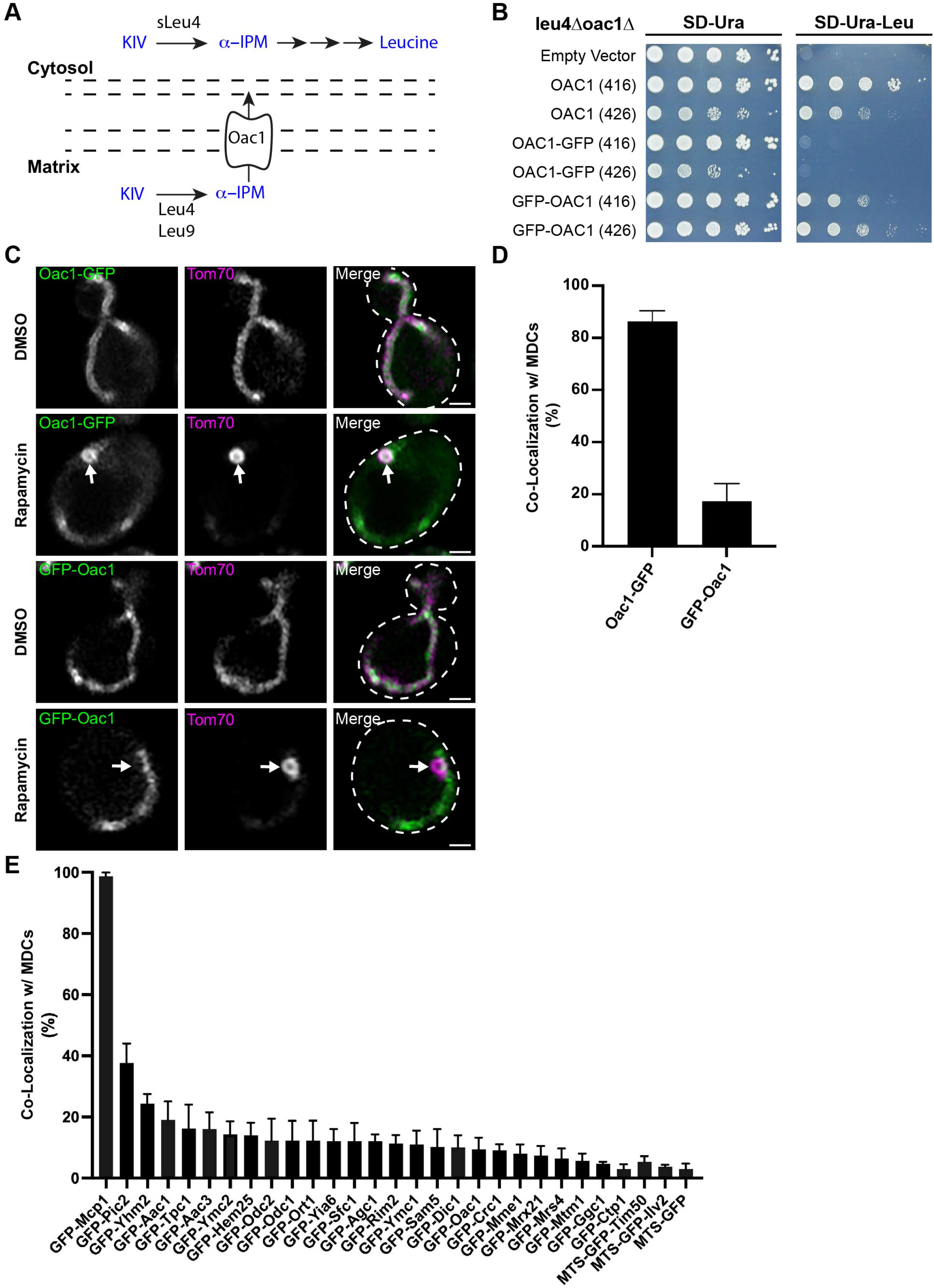
N-terminally tagged mitochondrial metabolite carriers are excluded from MDCs. (A) Diagram of Oac1 transport function within leucine biosynthesis in yeast. KIV, α- ketoisovalerate; α-IPM, α-isopropylmalate; sLeu4, cytosolic version of Leu4. (B) Cell growth spot assays of *leu4*Δ*oac1*Δ yeast transformed with the indicated Oac1 constructs expressed from either low-copy (pRS416) or high-copy (pRS426) plasmids, serially-diluted and plated on SD agar plates lacking either uracil (SD-Ura) or uracil and leucine (SD-Ura-Leu). (C) Super-resolution confocal fluorescence microscopy images of yeast expressing Tom70-mCherry and Oac1 endogenously tagged with GFP at the C-or N-terminus. Cells were treated with DMSO (vehicle control) or 200 nM rapamycin. The formation of MDCs in rapamycin treated cells are indicated by white arrows. Scale Bar= 1 µm (D) Quantification of the MDC co-localization frequency of the indicated Oac1 proteins in rapamycin treated cells. Error bars show mean ± standard error of three replicates, *n* ≥ 100 cells per replicate. (E) Quantification of the frequency N-terminally GFP-tagged mitochondrial metabolite carriers co-localize with MDCs. Error bars show mean ± standard error of three replicates, *n* ≥ 100 cells per replicate.

To determine if other mitochondrial metabolite carriers follow a similar pattern as that observed for Oac1, we analyzed whether mitochondrial metabolite carriers are still sequestered into MDCs when fused to GFP at their N-terminus. To do so, we examined strains from the N-terminal SWAp-Tag (N-SWAT) library (Weill et al., 2018), in which each metabolite carrier is fused at the N-terminus to superfolder GFP (sfGFP) and expressed under the control of the *NOP1* promoter. Indeed, the majority of N-terminally sfGFP-tagged metabolite carriers are excluded from entering MDCs unlike the C-terminally GFP-tagged carriers from the yeast GFP collection (Fig. 1 E and Fig. S1 A), which were originally analyzed in our fluorescence microscopy-based screen used to determine proteins incorporated into MDCs (Hughes et al., 2016). Importantly, sfGFP-Mcp1, an OMM protein previously identified to enrich in MDCs (Hughes et al., 2016) still strongly incorporated into MDCs in 100% of cells analyzed (Fig. 1 E and Fig. S1 A). Furthermore, Tim50 and Ilv2, two proteins excluded from MDCs when C-terminally fused to GFP (Schuler et al., 2021), were still excluded from MDCs when analyzed from the N-SWAT library (Fig. 1 E and Fig. S1 A). Thus, in contrast to our previous analyses on C-terminally tagged mitochondrial carriers (Hughes et al., 2016; Schuler et al., 2021), mitochondrial metabolite carriers are largely excluded from MDCs when visualized using an N-terminal GFP fusion.

### Cox7 and Cox8 differ in ability to drive cargo into MDCs

Other than mitochondrial metabolite carriers, the only other IMM-localized protein we previously identified in MDCs is Cox7, a subunit of cytochrome *c* oxidase (Fig. 2 A; Calder and McEwan 1991; Hughes et al., 2016). However, unlike Oac1, we found that the C-terminal GFP-tagged Cox7 was able to rescue growth of a *cox7*τ1 strain on non-fermentable carbon sources (Fig. S2 A), suggesting the protein is functional. Because our observations that the targeting of mitochondrial metabolite carriers to MDCs was altered by the location of the GFP epitope tag (Fig. 1 C-E), we conducted additional experiments to understand the parameters of Cox7 incorporation into MDCs. We assessed if Cox7 can dominantly drive a protein into MDCs through the creation of a chimera between Cox7 and the intermembrane space domain of Tim50. Tim50 is an essential component of the pre-sequence translocase of the inner membrane (Tim23) complex, and a protein strongly excluded from MDCs (Fig. 2 A; Hughes et al., 2016; Schuler et al., 2021). Prior work has demonstrated that the N-terminal portion (1-131) of Tim50, including its transmembrane domain, can be removed or replaced as long as the intermembrane space domain (132- 476) of Tim50 is maintained, as it provides the essential function of Tim50 (Mokranjac et al., 2009). Thus, we replaced the N-terminal portion of Tim50 with either full-length Cox7 or another subunit of cytochrome *c* oxidase, full-length Cox8 (Fig. 2 B). Both Cox7 and Cox8 are small (∼50 amino acids) proteins containing short matrix-localized segments at their N-terminus and a single C-terminal transmembrane domain (Calder and McEwan 1991; Patterson and Poyton, 1986). We analyzed the functionality of these Tim50 chimeras by assessing if they could rescue the essentiality of Tim50 through a plasmid shuffle growth assay. A *tim50*Δ yeast strain expressing Tim50 from a low-copy plasmid carrying a *URA3* selection marker was selected against with 5-fluorooroctic acid (5-FOA) and cell growth was maintained when an additional low-copy plasmid was present that expressed a functional Tim50-GFP protein (Fig. 2 C). Using this assay, we determined that a Cox8-Tim50-GFP chimera maintains the growth of a *tim50*Δ strain, while the Cox7-Tim50-GFP chimera failed to rescue the essentiality of Tim50 (Fig. 2 C), demonstrating that the Cox7-Tim50-GFP chimera is non-functional. Because all of these chimeric proteins still localize to mitochondria, we assessed if any these proteins are sequestered into MDCs. Notably, Cox7-Tim50-GFP strongly incorporated into MDCs, while both Tim50-GFP and Cox8-Tim50-GFP were excluded (Fig. 2, D and E). A primary difference between Cox7 and Cox8 is that Cox8 contains a well-annotated, N-terminal mitochondrial targeting presequence (MTS) (Vögtle et al., 2009). When we attached the MTS from Cox8 (codons 1-27) to the N-terminus of Cox7-Tim50-GFP (creating MTS-Cox7-Tim50-GFP; Fig. 2 B) we created a chimeric protein that could rescue the growth of *tim50*Δ cells (Fig. 2 C) and this chimeric protein was also excluded from MDCs (Fig. 2, D and E). Altogether, these results suggest that a key difference in the ability of Cox8 compared to Cox7 to support Tim50 functionality and MDC exclusion is the presence of an MTS.

**Figure 2.**
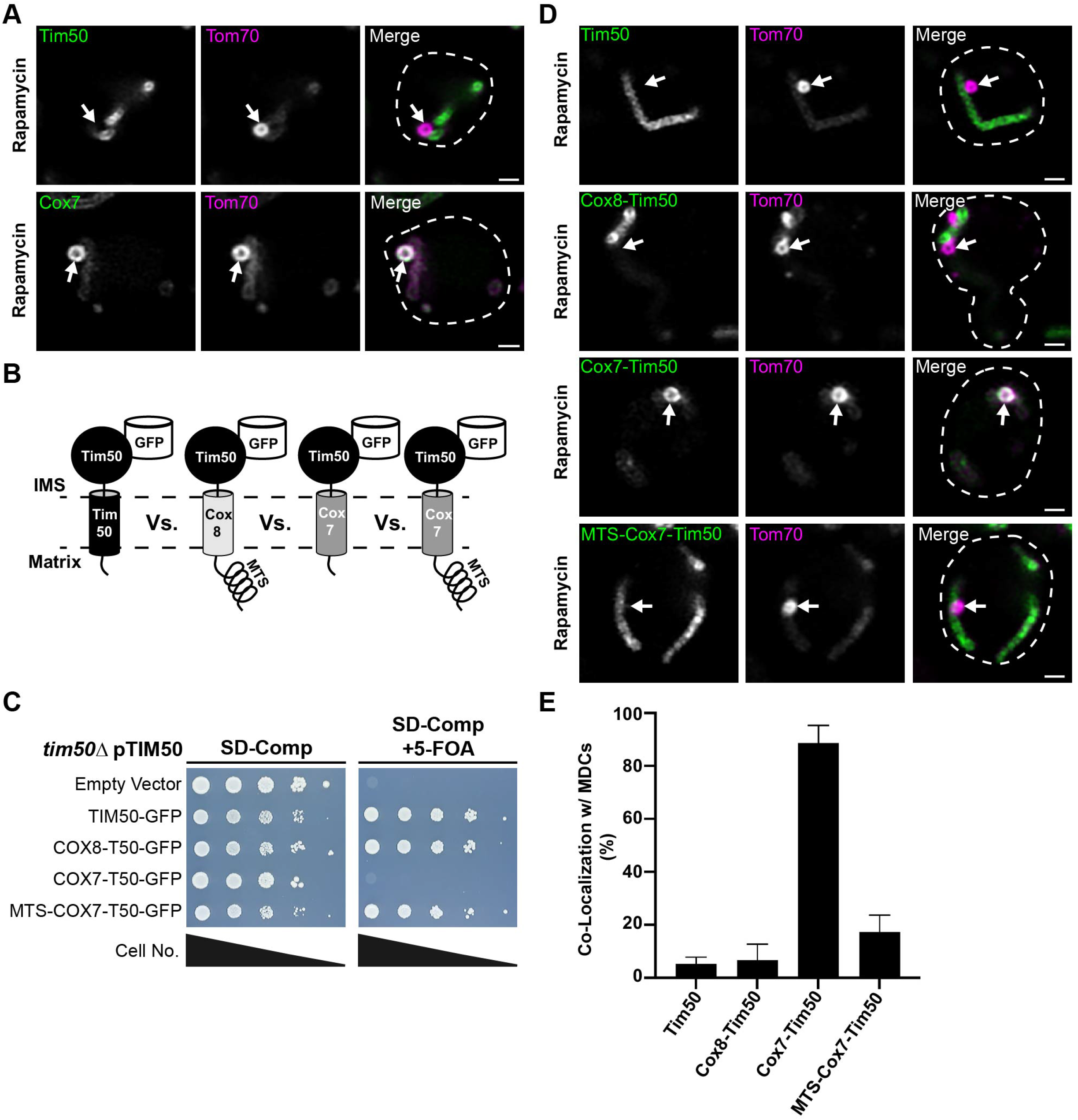
Cox7 and Cox8 differ in ability to drive Tim50 into MDCs. (A) Super-resolution confocal fluorescence microscopy images of rapamycin-induced MDC formation in yeast expressing Tom70-mCherry and Tim50-GFP or Cox7-GFP. MDCs are indicated by white arrows. Scale Bar= 1 µm (B) Schematic of the different Tim50 chimeras analyzed in C-E. (C) Cell growth spot assays of *tim50*Δ yeast transformed with a low-copy (pRS416) plasmid expressing Tim50 and additionally transformed with low-copy (pRS413) plasmids that express the indicated Tim50 chimeras. Cells were serially-diluted and plated on SD agar plates with a full complement of nutrients SD-Complete (SD-Comp) or SD-Complete with 1 µM 5-Fluoroorotic acid (SD-Comp + 5-FOA) (D) Super-resolution confocal fluorescence microscopy images of rapamycin-induced MDC formation in yeast expressing Tom70-mCherry and the indicated Tim50 chimeras. MDCs are indicated by the white arrows. Scale Bar= 1 µm (E) Quantification of the frequency the indicated Tim50 chimeras co-localize with MDCs. Error bars show mean ± standard error of three replicates, *n* ≥ 100 cells per replicate.

### The mitochondrial targeting sequence precludes cargo from entering MDCs

Considering that the addition of an MTS to Cox7-Tim50-GFP excluded this protein from entering MDCs, we examined if the addition of an MTS generally precludes proteins from MDC sequestration. We attached the MTS of Cox8 to Cox7-GFP, Oac1-GFP, and Tom70-GFP and assessed the co-localization of these proteins with MDCs. Indeed, while Cox7-GFP, Oac1-GFP, and Tom70-GFP are all strongly enriched within MDCs (consistent with our previous observations, Hughes et al., 2016; Schuler et al., 2021), the addition of an MTS to each of these proteins precluded them from entering MDCs (Fig. 3, A and B). Notably, adding an MTS to Cox7-GFP still produces a functional protein because it can rescue the growth of a *cox7*Δ yeast strain on medium containing a non-fermentable carbon source (Fig. S2 A). However, adding an MTS to Oac1-GFP or Tom70- GFP creates non-functional proteins (Fig. S2 B and C), which would be expected as the addition of an MTS to Oac1 would disrupt the orientation of Oac1 membrane insertion and MTS addition to Tom70 would be expected to drive the inward import of Tom70-GFP into mitochondria. Because mitochondrial targeting presequences facilitate the import of proteins into mitochondria, these results suggest that the subdomain localization of mitochondrial proteins is a key determinant for MDC sequestration.

**Figure 3.**
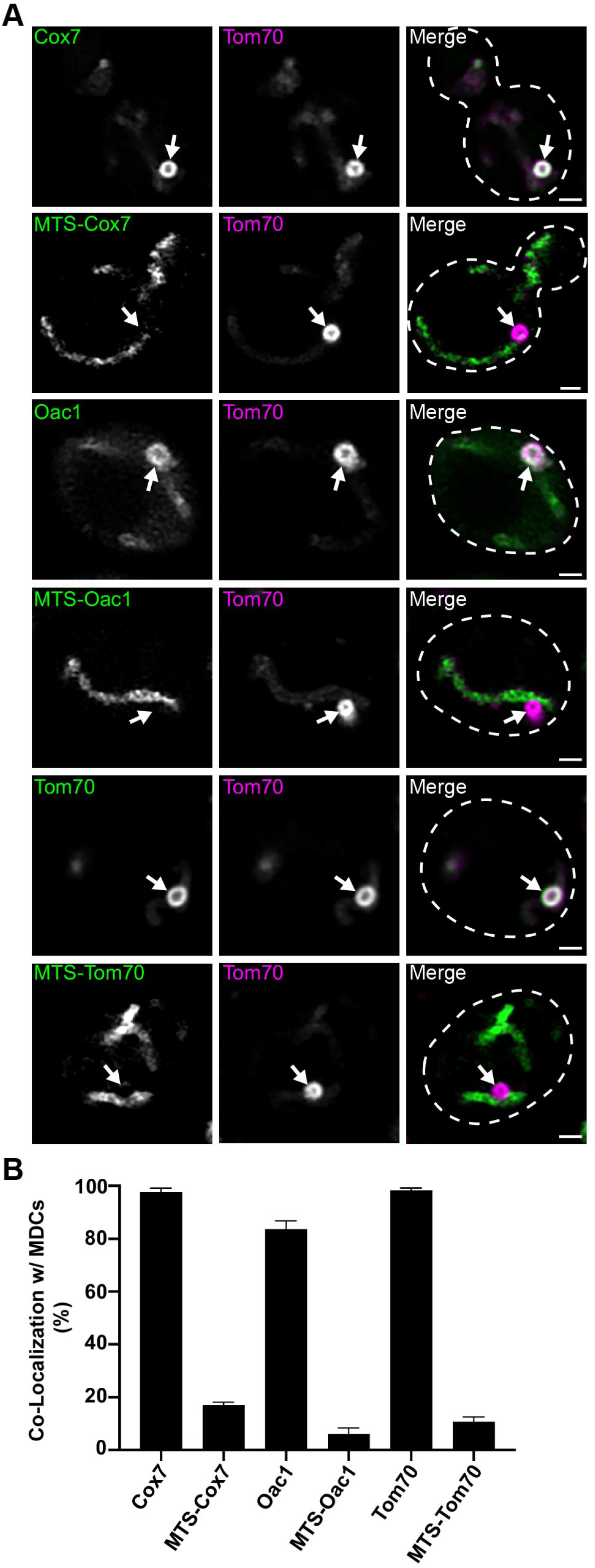
A mitochondrial targeting sequence precludes cargo from entering MDCs. (A) Super-resolution confocal fluorescence microscopy images of rapamycin-induced MDC formation in yeast expressing Tom70-mCherry and either Cox7-GFP, Oac1-GFP, or Tom70-GFP without or with an N-terminal mitochondrial targeting sequence (MTS). MDCs are indicated by the white arrows. Scale Bar= 1 µm (B) Quantification of the frequency the indicated proteins co-localize with MDCs. Error bars show mean ± standard error of three replicates, *n* ≥ 100 cells per replicate.

### MDC cargo are localized to the mitochondrial outer membrane

To assess the mitochondrial subdomain localization of proteins differentially incorporated into MDCs, we initially used a protease protection assay. While our results supported the interpretation that Cox7-GFP is more restricted to the OMM than MTS-Cox7-GFP and not clearly localized in the IMM, we also observed that some of the membrane proteins we investigated formed protease resistant populations, including Cox7-GFP, complicating our interpretations of the results from this protease-based assay (Fig. S3 A, see below for MTS-Oac1-GFP). Instead, we adopted a different strategy based on the observation that we could directly visualize isolated mitochondria using super-resolution confocal fluorescence microscopy to assess protein localization within mitochondria. Using this strategy, we could resolve that Tim50-GFP was internally located within mitochondria compared to Tom70-mCherry, an outer membrane protein (Fig. 4 A). Similarly, both Cox15-GFP (inner membrane protein) and Lat1-GFP (matrix protein) showed a diffuse internal fluorescence inside mitochondria compared to Tom70-mCherry (Fig. 4 A). Alternatively, mitochondria isolated from cells expressing Tom20-GFP and Tom70- mCherry showed that these proteins co-localized, which would be expected as both are OMM proteins (Fig. 4 A). These results are consistent with prior observations on the mitochondrial subdomain localizations for each of these proteins (Vögtle et al., 2017) and demonstrates that direct visualization of isolated mitochondria *in vitro* can be utilized to identify proteins localized to the OMM from those localized to the interior of mitochondria.

**Figure 4.**
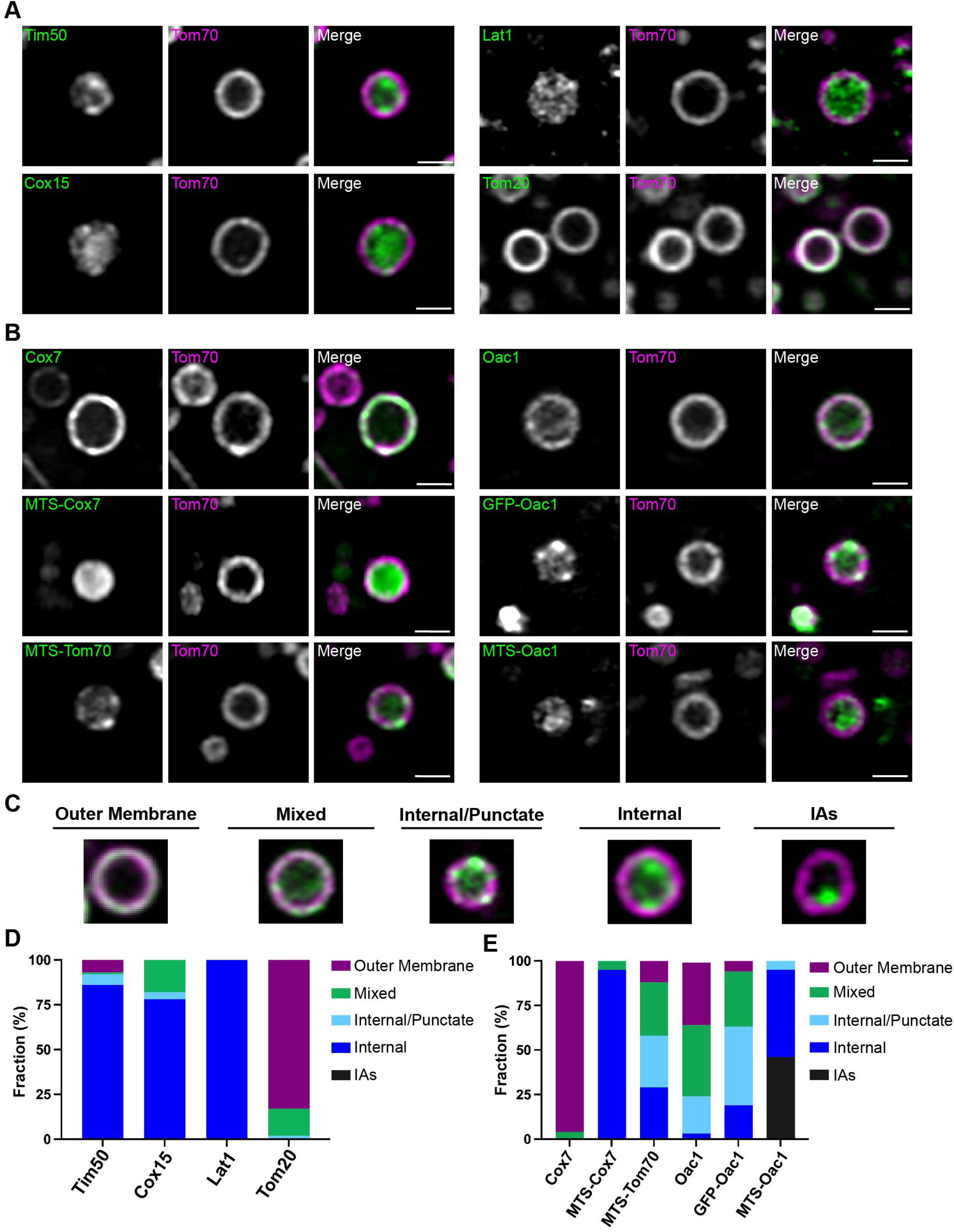
MDC cargo are localized to the outer mitochondrial membrane. (A) Super-resolution confocal fluorescence microscopy images of purified mitochondria from cells expressing Tom70-mCherry and either Tim50-GFP, Cox15-GFP, Lat1-GFP, or Tom20-GFP. Scale Bar= 1 µm (B) Super-resolution confocal fluorescence microscopy images of purified mitochondria from cells expressing Tom70-mCherry and either Cox7-GFP, MTS-Cox7-GFP, MTS-Tom70-GFP, Oac1-GFP, GFP-Oac1, or MTS-Oac1-GFP. Scale Bar= 1 µm (C) Examples of the diverse protein localizations observed in purified mitochondria compared to Tom70-mCherry used for the quantifications in D and E. (D) Quantification of the localization of each indicated protein imaged in A within purified mitochondria shown as a percent of the total mitochondria observed, *n* ≥ 100. (E) Quantification of the localization of each indicated protein imaged in B within purified mitochondria shown as a percent of the total mitochondria observed, *n* ≥ 100.

Occasionally, we found that protein localizations were more complex than either overlapping with Tom70-mCherry or residing inside the Tom70-mCherry marked outer membrane ring (Fig. 4 B). To quantify our observations, we assigned protein localizations to the following categories: overlapping with Tom70-mCherry (outer membrane), overlapping with Tom70-mCherry and internal luminal fluorescence (mixed), internal luminal fluorescence with discrete puncta that overlapped with Tom70-mCherry (internal/punctate), internal luminal fluorescence only (internal), and bright internal puncta (internal aggregates, IAs) (Fig. 4 C). Using these categories to quantify protein localizations for several of our control proteins, we observed that Tim50-GFP is internally localized in ∼80% of isolated mitochondria, Cox15-GFP is internally localized in ∼75% of mitochondria, and Lat1-GFP was always internally localized, while Tom20-GFP strongly overlapped with Tom70-mCherry on the outer membrane of isolated mitochondria (Fig. 4 D). These results are consistent with observations performed *in vivo* using swollen mitochondria to determine the subdomain localizations of mitochondrial proteins (Wurm and Jakobs, 2006), except they can additionally resolve intramitochondrial proteins that localize to the inner boundary membrane, such as Tim50.

Using this fluorescence microscopy strategy to analyze protein localizations on isolated mitochondria, we observed that both Cox7-GFP and Oac1-GFP, two proteins that incorporate into MDCs, strongly co-localized with Tom70-mCherry on the outer membrane of isolated mitochondria (Fig. 4, B and E). Conversely, MTS-Cox7-GFP and MTS-Oac1-GFP, two protein constructs that are excluded from MDCs, were strongly localized within the interior of isolated mitochondria (Fig. 4, B and E). Note that MTS-Oac1-GFP frequently formed internal aggregates (Fig. 4, B and E), which was not surprising considering that the addition of an MTS to Oac1 would be expected to disrupt the topology of Oac1 within the IMM. Both GFP-Oac1 and MTS-Tom70-GFP, two other constructs that were excluded from MDCs, showed more of an internal localization within isolated mitochondria (Fig. 4, B and E), albeit not as robustly as MTS-Cox7-GFP or MTS-Oac1-GFP. We also observed a similar intramitochondrial localization for MTS-Cox7-GFP and MTS-Oac1-GFP by acutely swelling mitochondria *in vivo* (Fig. S3 B). However, examining swollen mitochondria *in vivo* it is not possible to distinguish some proteins (Tim50 and Oac1) expected to localize to the inner boundary membrane from proteins (Tom70) that localize to the OMM (Fig. S3 B, Wurm and Jakobs, 2006). One additional possibility for the difference in protein localization of the MTS-containing membrane proteins is that they were aberrantly imported into the mitochondrial matrix. However, upon using a carbonate extraction assay to examine the integration of these proteins within mitochondrial membranes, we observed that MTS-Oac1-GFP, MTS-Cox7-GFP, and MTS-Tom70-GFP all integrate into membranes to a similar extent as the non-MTS containing versions (Fig. S3, C-E). In total, these mitochondrial subdomain localization studies demonstrate that the majority of Cox7-GFP and Oac1-GFP proteins are aberrantly localized to the OMM and the subdomain localization of proteins within mitochondria is a key determinant for their sequestration into MDCs.

Previously, we determined that all detectable mitochondrial matrix proteins and IMM proteins (which now include mitochondrial metabolite carriers and Cox7) are excluded from MDCs, and originally annotated that IMS proteins are also excluded from MDCs (Hughes et al., 2016). We considered that IMS proteins may still enter MDCs but potentially could not be detected in our initial widefield fluorescence microscopy-based screen. However, using super-resolution confocal fluorescence microscopy we still observed that several IMS proteins are strongly excluded from MDCs (Fig. S3 F). Thus, our results suggest that MDCs incorporate membrane cargo proteins exclusively from the OMM and restrict proteins from intramitochondrial subdomains.

### Subunits of the TOM complex are excluded from MDCs unless complex assembly is impaired

While MDC cargo are incorporated from the OMM, we have also observed that not all OMM proteins are sequestered into MDCs to the same extent. For example, the OMM receptor for MTS-containing mitochondrial proteins, Tom20, is incorporated into MDCs, but unlike Tom70, it is not enriched within MDCs (Fig. 5 A; Schuler et al., 2021). A line-scan analysis showed that the amount of Tom20-GFP incorporated into MDCs is similar to the concentration of Tom20-GFP found throughout the mitochondrial tubule, highlighted by a mean MDC to mitochondrial tubule fluorescence intensity ratio close to 1 (1.22; Fig. 5, A and B). This observation is in contrast to Tom70-mCherry, which is strongly enriched within MDCs with a mean MDC to mitochondrial tubule fluorescence intensity ratio of 8.20 (Fig. 5 B). To begin to assess how Tom20 is limited from MDC sequestration, we truncated Tom20 at regions flanking the tetratricopeptide motif previously shown to bind MTS-containing precursor proteins (Abe et al., 2000), creating a truncation that only maintained the N-terminal transmembrane domain of Tom20 (Tom20^1-50^-GFP) and a truncation that only removed the final 28 C-terminal amino acids of Tom20 (Tom20^1-145^-GFP). Intriguingly, while Tom20^1-145^-GFP acted similar to full-length Tom20-GFP and was not enriched within MDCs (Fig. 5, A and B), Tom20^1-50^-GFP was robustly sequestered within MDCs (Fig. 5 A), highlighted by a 3-fold enrichment into MDCs (Fig. 5 B), demonstrating that the Tom20 C-terminal precursor-binding domain is required to limit Tom20 incorporation into MDCs. Notably, we also observed that removing the precursor-binding domain of Tom70 (Tom70^1-98^-GFP) did not disrupt the robust accumulation of this protein into MDCs (Fig. 5, A and B), but appeared to increase enrichment within MDCs, potentially indicating that an OMM-targeting transmembrane domain, by itself, is sufficient for MDC sequestration.

**Figure 5.**
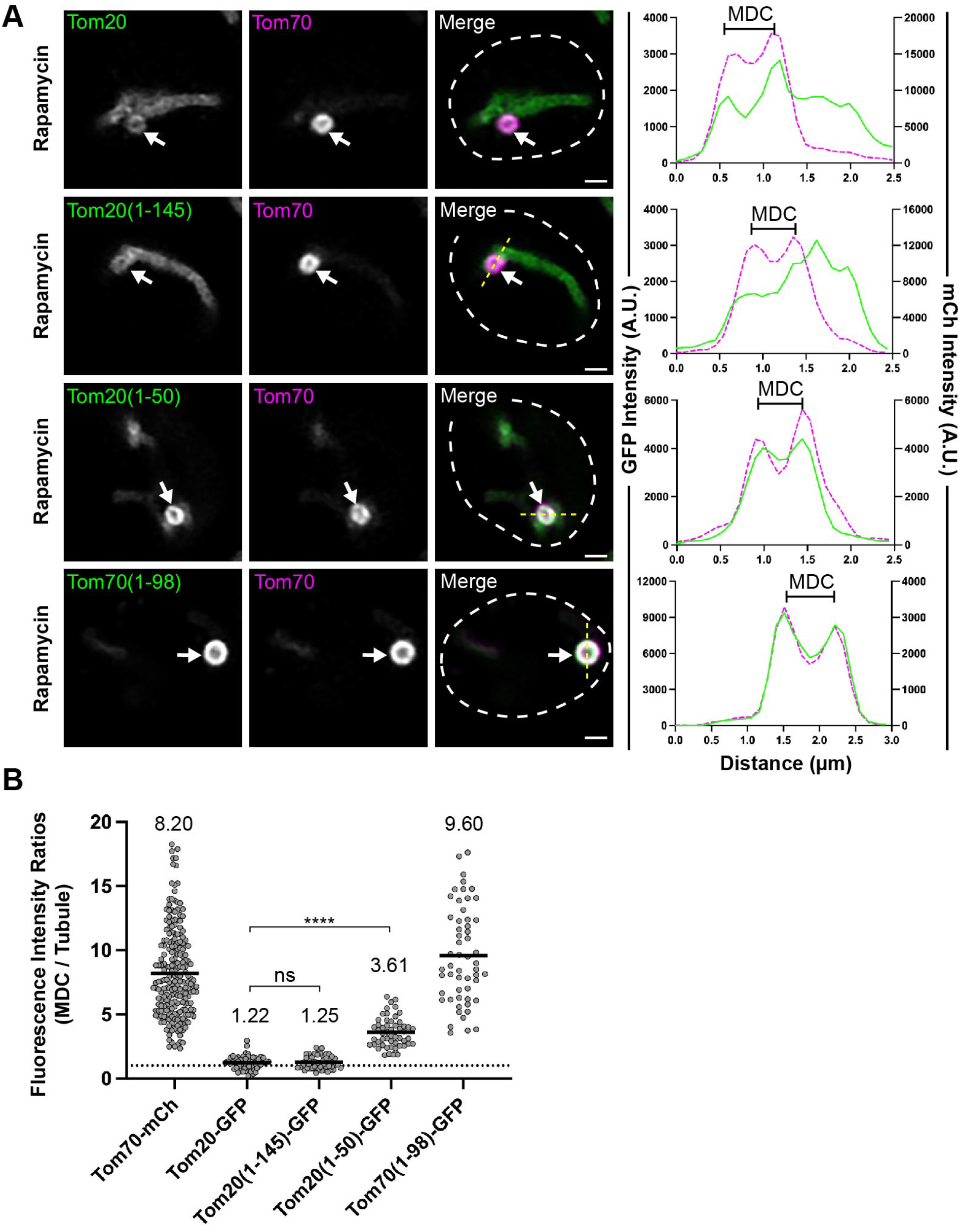
Tom20 becomes enriched in MDCs when truncated to its N-terminal transmembrane domain. (A) Super-resolution confocal fluorescence microscopy images of rapamycin-induced MDC formation in yeast expressing Tom70-mCherry and the indicated Tom20-GFP proteins or a truncated version of Tom70, Tom70^1-98^-GFP. MDCs are indicated by the white arrows. Scale Bar= 1 µm. Yellow line marks the position of the line-scan fluorescence intensity profile. Left and right Y axis correspond to the GFP and mCherry fluorescence intensity, respectively. Bracket marks MDC. (B) Quantification of protein enrichment within MDCs from the yeast strains analyzed in A. Each dot represents the ratio of the fluorescence intensity measured within MDCs compared to the fluorescence intensity in the mitochondrial tubule. Dark line indicates the mean with the mean value shown above. The dotted line demarcates a ratio of 1. Unpaired *t* tests with Welch’s correction were used to determine statistical significance; ns, not significant; ****, *P* < 0.0001.

Prompted by our results analyzing Tom20, we were curious if core subunits of the translocase of the outer membrane (TOM) complex also showed a limited enrichment within MDCs. Strikingly, GFP-Tom22, GFP-Tom5, and GFP-Tom7 all showed a strong de-enrichment within MDCs, as each of these TOM complex subunits were observed to have mean MDC to mitochondrial tubule fluorescence intensity ratios below 1 (Fig. 6, A-D). These results further highlight that there are additional selection criteria for incorporating proteins within MDCs than just being localized to the OMM. To assess if assembly into TOM complexes is a key determinant restricting these subunits from entering MDCs we analyzed MDC enrichment of TOM complex subunits in cells lacking *TOM6* (*tom6*Δ), which encodes a key but non-essential subunit of the TOM complex that supports complex assembly (Dekker et al., 1998). Intriguingly, within *tom6*Δ cells, GFP-Tom22, GFP-Tom5, and GFP-Tom7 were all robustly sequestered within MDCs with a comparable 3-fold enrichment within MDCs (Fig. 6, A-D), demonstrating that core subunits of the TOM complex are typically precluded from MDC sequestration unless TOM complex assembly is impaired.

**Figure 6.**
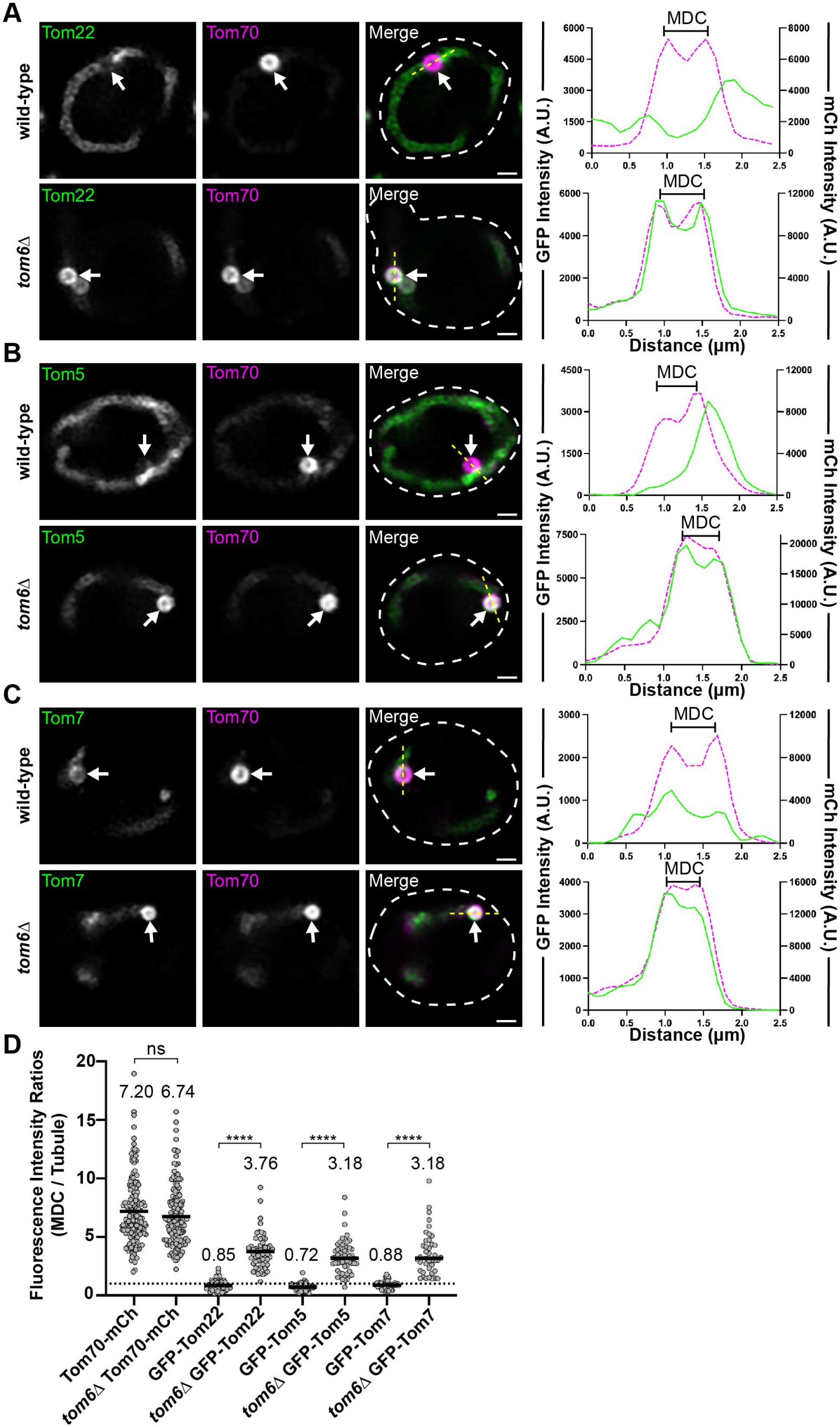
Subunits of the TOM complex are excluded from MDCs unless complex assembly is impaired. (A-C) Super-resolution confocal fluorescence microscopy images of rapamycin-induced MDC formation in wild-type or *tom6*Δ yeast expressing Tom70-mCherry and the indicated TOM complex subunits N-terminally tagged with GFP. MDCs are indicated by the white arrows. Scale Bar= 1 µm. Yellow line marks the position of the line-scan fluorescence intensity profile. Left and right Y axis correspond to the GFP and mCherry fluorescence intensity, respectively. Bracket marks MDC. (D) Quantification of protein enrichment within MDCs from wild-type or *tom6*Δ yeast strains analyzed in A-C. Each dot represents the ratio of the fluorescence intensity measured within MDCs compared to the fluorescence intensity in the mitochondrial tubule. Dark line indicates the mean with the mean value shown above. The dotted line demarcates a ratio of 1. Unpaired *t* tests with Welch’s correction were used to determine statistical significance; ns, not significant; ****, *P* < 0.0001.

### Overloading the OMM with membrane proteins induces MDC formation

An analysis of MDC formation demonstrated that OMM-derived extensions repeatedly elongate, coalesce, and invaginate to generate compartments that capture OMM cargo and cytosol within a protected domain (Wilson et al., 2023 *preprint*). These structures are similar to the membrane-enriched domains generated by the ER in response to the overexpression and aberrant accumulation of some ER membrane proteins, which has been described as a form of ER microautophagy (Wright et al., 1988; Schuck et al., 2014; Schäfer et al., 2020). Previously, we observed that overexpression of Tom70 or several mitochondrial metabolite carriers induces constitutive MDC formation and also sensitizes cells to produce more MDCs in response to Concanamycin A treatment, a specific inhibitor of the Vacuole H^+^-ATPase (Schuler et al., 2021, see also Fig. 7A). Additionally, in a separate study we noticed that constitutive overexpression of the OMM protein Mcp1 in wild-type yeast also induced constitutive MDC formation (Fig. 7 A; English et al., 2020). Due to these results and the striking similarity between MDCs and ER microautophagy, we examined if the overexpression of OMM proteins similarly induce the generation of MDCs from mitochondria. Indeed, a vast array of OMM proteins were all capable of inducing constitutive MDC formation when individually overexpressed (Fig. 7 A and Figure S4 A). Notably, all of the proteins that led to robust MDC formation, with the exception of Tom7, are all OMM proteins that are strongly sequestered in MDCs (Fig. S4 B). Additionally, the particular OMM proteins that elicited a robust MDC formation response all had diverse membrane topologies and are embedded in the OMM through diverse translocation pathways (Wiedemann and Pfanner, 2017). Because Tom70 facilitates the targeting and import of OMM proteins and performs an unknown supportive role in MDC formation (Hughes et al., 2016), we assessed if Tom70 is required for MDCs to form in response to the overexpression of OMM proteins. In *tom70*Δ cells, we analyzed MDC formation upon the constitutive overexpression of ten OMM proteins or the metabolite carrier, Oac1, all of which had produced the strongest MDC formation in wild-type cells. In most cases, we observed that MDC formation was strongly blunted in *tom70*Δ cells, with the exception of Scm4 and Pth2, both of which still induced MDCs to form in ∼20% of cells (Fig. S4 C).

**Figure 7.**
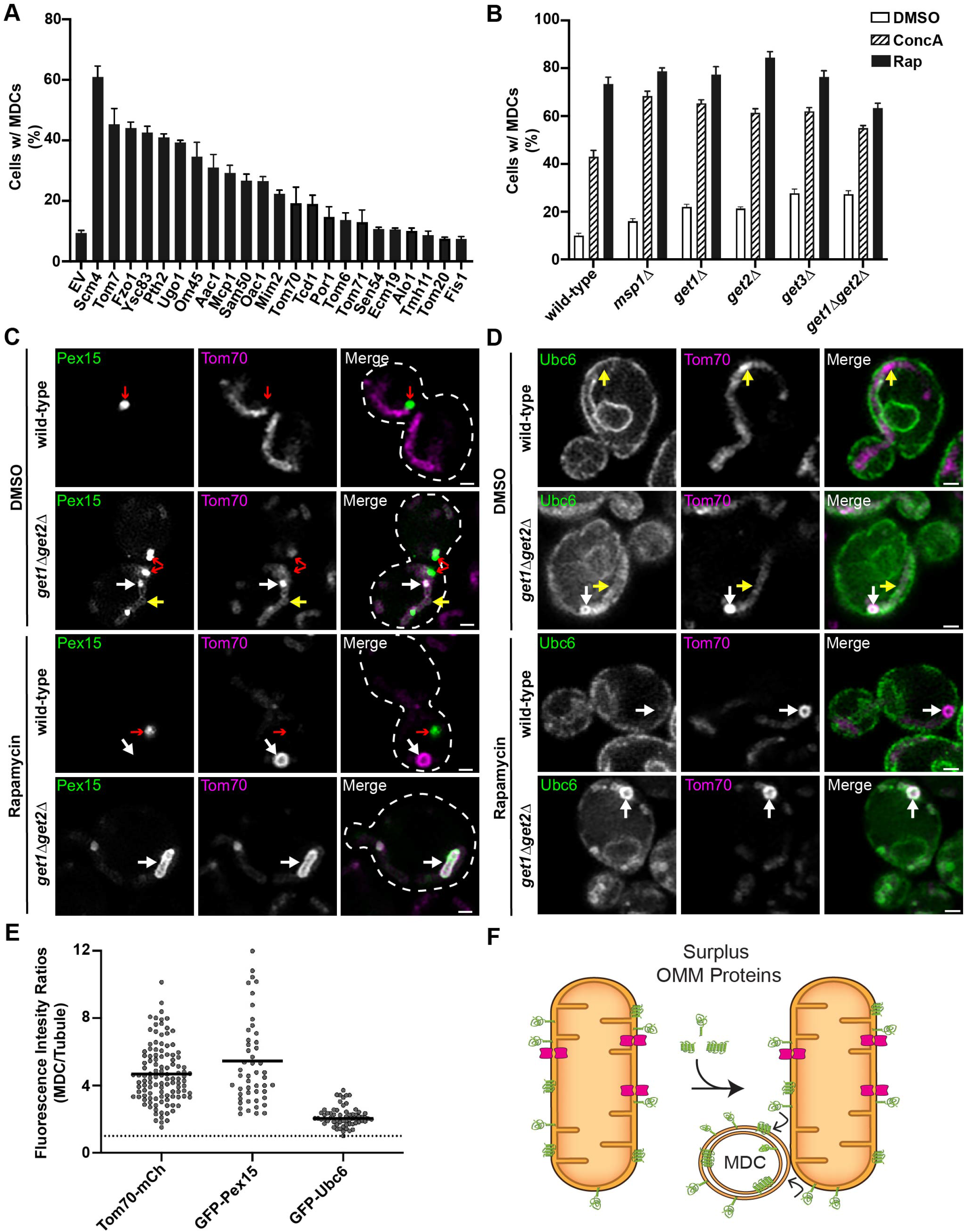
Overloading the OMM with membrane proteins induces MDC formation. (A) Quantification of the percent of cells forming MDCs in cells constitutively overexpressing the indicated proteins. Error bars show mean ± standard error of three replicates, *n* ≥ 100 cells per replicate. (B) Quantification of the percent of cells forming MDCs in the indicated yeast strains treated with DMSO (vehicle control), 500 nM Concanamycin A (ConcA), or 200 nM Rapamycin (Rap). Error bars show mean ± standard error of three replicates, *n* ≥ 100 cells per replicate. (C) Super-resolution confocal fluorescence microscopy images of wild-type or *get1*Δ*get2*Δ yeast strains expressing Tom70-mCherry and overexpressing GFP-Pex15 treated with DMSO or 200 nM rapamycin. Yellow arrows indicate mitochondrial tubules, white arrows indicate MDCs, red arrows indicate peroxisomes. Scale Bar= 1 µm. (D) Super-resolution confocal fluorescence microscopy images of wild-type or *get1*Δ*get2*Δ yeast strains expressing Tom70-mCherry and overexpressing GFP-Ubc6 treated with DMSO or 200 nM rapamycin. Yellow arrows indicate mitochondrial tubules, white arrows indicate MDCs. Scale Bar= 1 µm. (E) Quantification of protein enrichment within MDCs from the yeast strains analyzed in C and D. Each dot represents the ratio of the fluorescence intensity measured within MDCs compared to the fluorescence intensity in the mitochondrial tubule. Dark line indicates the mean with the mean value shown above. The dotted line demarcates a ratio of 1. (F) Model of MDC sequestration of surplus proteins from the OMM.

As our results suggest that MDCs can sequester and remove excess proteins from the OMM, we hypothesized that the MDC pathway functions in concordance with other mitochondrial quality control pathways known to also remove excess OMM proteins. To investigate this idea, we analyzed MDC formation in *msp1*Δ cells, which lack the mitochondrial membrane-embedded AAA-ATPase shown to extract mistargeted tail-anchored (TA) OMM proteins (Chen et al., 2014). We observed that *msp1*Δ cells had both an increase in steady-state MDC formation and enhanced MDC formation after treatment with ConcA and a modest increase upon rapamycin treatment (Fig. 7 B). Consistent with these results, we also observed enhanced MDC formation in *get1*Δ, *get2*Δ, *get3*Δ, and *get1*Δ*get2*Δ cells, which lack key proteins required for TA protein targeting and insertion at the ER, leading to enhanced mistargeting of TA proteins to the OMM (Fig. 7 B; Schuldiner et al., 2008). Prompted by these results, we examined if disrupting other ER-protein import pathways also led to enhanced MDC formation. To do so, we generated genetic deletions to remove non-essential subunits of the Sec63 complex (*sec72*Δ), Ssh1 translocon complex (*ssh1*Δ), SRP-independent protein targeting (*snd3*Δ), and the ER transmembrane complex (*emc2*Δ) and analyzed MDC formation (Green et al., 1992; Finke et al., 1996; Aviram et al., 2016; Jonikas et al., 2009). While *ssh1*Δ cells demonstrated a dramatic increase in constitutive MDC formation and enhanced MDC formation upon ConcA treatment, disruption of the other ER targeting and import pathways did not significantly affect MDC formation (Fig. S4 D).

Because MDCs strongly sequester some but not all OMM proteins, we were curious if MDCs were capable of sequestering mistargeted TA proteins. When we ectopically overexpressed GFP-Ubc6 and GFP-Pex15, two TA proteins that mistarget to the OMM in cells with an impaired GET pathway (Schuldiner et al., 2008; Chen et al., 2014), we observed that both GFP-Ubc6 and GFP-Pex15 localized more frequently to mitochondria in *get1*Δ*get2*Δ yeast compared to wild-type yeast (yellow arrows, Fig. 7 C and D; Fig. S4 E). Consistent with our results that MDCs form more frequently in *get1*Δ*get2*Δ cells, we also observed the enrichment of GFP-Pex15 and GFP-Ubc6 within MDCs in vehicle-treated cells (white arrows, Fig. 7 D). Upon rapamycin treatment, mitochondrial-localized GFP-Pex15 and GFP-Ubc6 were always observed to co-localize with MDCs (Fig. 7, C and D), and these proteins were also enriched within MDCs (Fig. 7 E), demonstrating that MDCs are capable of sequestering mistargeted proteins. Collectively, we observed that MDCs preferentially ensnare membrane protein cargo from a single membrane, the OMM, that MDCs form in response to disturbances in OMM protein concentration, and that through the formation of an MDC both membrane and membrane protein cargoes are sequestered into a distinct, protected domain (Fig. 7 F).

## DISCUSSION

The outer mitochondrial membrane creates a critical interface between mitochondria and the rest of the cell. While the OMM supports mitochondrial biogenesis and establishes contacts with other organelles, it can also be a target for both cellular proteins mistargeted to the mitochondrial surface and pathogen proteins (Schuldiner et al., 2008; Costa et al., 2018; Li et al., 2022). In response, mitochondria have several mechanisms to safeguard the OMM, including the remodeling of the OMM to package aberrant proteins into distinct domains, which can subsequently be removed and degraded by lysosomes (Soubannier et al., 2012; Hughes et al., 2016). Previously, we identified a piecemeal autophagic mechanism, called mitochondrial-derived compartments (MDCs), that robustly sequesters some mitochondrial membrane proteins and delivers them to yeast vacuoles for degradation (Hughes et al., 2016). Through an examination of protein selectivity into MDCs, we have shown here that MDCs concentrate cargo proteins from a single membrane, the OMM, and exclude proteins from internal mitochondrial domains. These results provide an explanation for how only a minor subset of mitochondrial proteins become robustly incorporated into MDCs. Based on these results and observations from an accompanying manuscript (Wilson et al., 2023 *preprint*) that demonstrates that MDCs form from OMM extensions that inwardly engulf layers of OMM, we considered that MDCs may act as a membrane-enriched trap to sequester surplus protein cargoes from the OMM. In support, we observed that the overexpression of OMM proteins, as well as, the mistargeting of TA proteins to the OMM all induced constitutive MDC formation. Furthermore, these aberrant proteins were sequestered into MDCs, which is not the case for all proteins residing in the OMM, as subunits of the TOM complex are excluded from MDCs, unless their assembly or association with the TOM complex becomes impaired. In total, these results suggest that MDCs may facilitate the removal of membrane and protein from mitochondria while maintaining critical functional aspects of the organelle.

Form is often indicative of function. In an accompanying manuscript we show that MDCs form through mitochondria creating OMM extensions that repeatedly elongate, coalesce, and invaginate to create a domain that not only concentrates OMM cargo, but can also segregate OMM cargo into a protected compartment (Wilson et al., 2023 *preprint*). Similar membrane proliferations and the formation of multilamellar whorls are generated from ER membranes in response to the accumulation of certain membrane proteins and in response to ER stress (Wright et al., 1988; Schuck et al., 2009; Schäfer et al., 2019). These ER membrane proliferations have been described as a form of ER microautophagy, as they are eventually directly engulfed by vacuoles/lysosomes (Schuck et al., 2014). Prompted by the similarities between ER microautophagy and MDC formation and because mitochondria utilize quality control pathways that share components with or directly engage with ER quality control pathways (Mårtensson et al., 2019; Matsumoto et al., 2019), we examined and observed that MDCs similarly arise from the accumulation of proteins at the OMM. Considering that MDCs also concentrate membrane, it seems likely that MDC formation may occur in response to a variety of mechanisms that challenge OMM integrity and thus could sequester diverse hydrophobic cargo. In support, bacteria also shed their outer membrane in response to both the presence of aberrant proteins and to alterations in membrane composition (Schwechheimer and Kuehn, 2013; Roier et al., 2016). In addition, we have observed OMM-enriched MDC structures in mammalian cell lines (Schuler et al., 2020 *preprint*), and mammalian mitochondria have been observed to shed their OMM in response to infection-induced stress through a structure that shares many characteristics with MDCs (Li et al., 2022). We have also found that alterations in cellular and mitochondrial lipid profiles occur during MDC formation induced by several acute stressors and that the two phospholipids synthesized in mitochondria, cardiolipin (CL) and phosphatidylethanolamine (PE), differentially affect MDC biogenesis (Xiao et al., 2023 *preprint*).

How cargo proteins are selectively targeted to MDCs still needs to be determined. We consistently observe a robust concentration of Tom70 within MDCs and MDC formation is impaired in the absence of Tom70 (Hughes et al., 2016; Schuler et al., 2021). Tom70 is a multifunctional protein that has been demonstrated to be involved in diverse mitochondrial functions, including protein import, mitochondrial biogenesis, organelle contacts, and protein quality control pathways (Söllner et al., 1990; Elbaz-Alon et al., 2015; Weidburg and Amon, 2018; Backes et al., 2021; Liu et al., 2022). Thus, Tom70 may perform various roles in supporting MDC formation. However, it is notable that a premier function of Tom70 is as a co-chaperone and receptor for hydrophobic precursor proteins and other aggregation prone proteins (Backes et al., 2018; Backes et al., 2021), many of which are MDC cargo proteins, including mitochondrial metabolite carriers when they become restricted to the mitochondrial surface. This function of Tom70 may indicate that Tom70 could act as a protein chaperone that additionally maneuvers cargo proteins into MDCs.

In the formation of an MDC, the OMM would need to proliferate without a connection to the IMM, and thus proteins that can more readily diffuse through the OMM may become preferentially captured by MDCs. Such a mechanism might provide a convenient method to capture aberrant proteins that do not have strong, restrictive interactions with other mitochondrial proteins. This possibility may explain why Tom70, a protein known to be diffusely spread throughout the OMM, is robustly sequestered into MDCs, while subunits of the TOM complex are de-enriched from MDCs, as TOM complexes localize to intramitochondrial contact sites to mediate protein import (Chacinska et al., 2010; Schulte et al., 2023). Furthermore, a diffusion mechanism may also explain why truncating both Tom20 and Tom70 to only their OMM-targeting N-terminal transmembrane domains was sufficient for the robust incorporation of these proteins into MDCs, and could explain how disrupting TOM complex assembly in the *tom6*Δ strain led to TOM complex subunits becoming sequestered into MDCs. It is also intriguing that we observed the exclusion of the TOM complex from MDCs, a result that suggests that MDCs preserve key processes involved in mitochondrial biogenesis and as such, could facilitate organelle remodeling in response to diverse stress or metabolic conditions.

Several quality control mechanisms have been identified that monitor the integrity of the OMM by removing precursor proteins from stalled protein import complexes or by removing mistargeted, misfolded, or damaged proteins from the OMM (Heo et al., 2010; Weidberg and Amon, 2018; Mårtensson et al., 2019, Metzger et al., 2020). Our results suggest that MDCs function in concordance with these pathways, but rather than directly extracting aberrant proteins from the OMM, the OMM is instead remodeled into a sequestering domain. Protein and membrane sequestration into MDCs may function as a distinct quality control mechanism that is induced when other OMM monitoring pathways become overwhelmed, which is supported by the enhanced MDC formation that occurs in cells lacking Msp1. However, we have also previously observed that Msp1 is another protein strongly sequestered into MDCs, along with Tom70, both of which function in several protein quality control pathways (Chen et al., 2014; Weidburg and Amon, 2018; Backes et al., 2021). Additionally, we have observed that MDCs make abundant contact with the ER (English et al., 2020; Wilson et al., 2023 *preprint*), an organelle that facilitates mitochondrial protein targeting (Hansen et al., 2018) and quality control (Matsumoto et al., 2019; Matsumoto et al., 2023). It seems possible that MDCs could provide a distinct membrane platform that concentrates aberrant proteins or precursor proteins and functions with other quality control mechanisms to efficiently remove proteins from the organelle.

## EXPERIMENTAL MODEL AND SUBJECT DETAILS

### Yeast Strains

All yeast strains are derivatives of *Saccharomyces cerevisiae* S288C (BY) (Brachmann et al., 1998) and are listed in Table S1. Deletion strains were created by one-step PCR-mediated gene replacement using the previously described pRS series of vectors (Brachmann et al., 1998; Sikorski and Hieter, 1989) and oligo pairs listed in Table S2. Correct gene deletions were confirmed by colony PCR across the chromosomal insertion site. Strains expressing proteins with attached C-terminal fluorescent proteins were created by one step PCR-mediated C-terminal endogenous epitope tagging using standard techniques and oligo pairs listed in Table S2. Plasmid templates for fluorescent epitope tagging were from the pKT series of vectors (Sheff and Thorn, 2004). Correct integrations were confirmed by a combination of colony PCR across the chromosomal insertion site and correctly localized expression of the fluorophore by microscopy. An integrated yEGFP-Oac1 fusion expressed from its endogenous promoter was generated by integrating a PCR construct containing the URA3 gene flanked by two halves of yEGFP directly upstream of the OAC1 ORF. URA3 was subsequently selected against by 5F-OA and the in-frame integration of yEGFP at the N-terminus of Oac1 was determined by sequencing. Where indicated, strains expressing proteins with superfolder GFP fused to the N-terminus are from the SWAp-Tag library described in Weill et al., 2018 and were a gift from Maya Schuldiner.

## METHOD DETAILS

### Yeast Cell Culture and Growth Assays

Yeast cells were grown exponentially for 15-16 hours at 30°C to a final optical (wavelength 600nm) density of 0.5 - 1 before the start of all experiments. This period of overnight log-phase growth was carried out to ensure vacuolar and mitochondrial uniformity across the cell population and is essential for consistent MDC formation. Unless otherwise indicated, cells were cultured in YPAD medium (1% yeast extract, 2% peptone, 0.005% adenine, 2% glucose). Otherwise, cells were cultured in a synthetic defined (SD) medium that contained the following unless specific nutrients were removed to select for growth or plasmid retention: 0.67% yeast nitrogen base without amino acids, 2% glucose, supplemented nutrients 0.072 g/L each adenine, alanine, arginine, asparagine, aspartic acid, cysteine, glutamic acid, glutamine, glycine, histidine, myo-inositol, isoleucine, lysine, methionine, phenylalanine, proline, serine, threonine, tryptophan, tyrosine, uracil, valine, 0.369g/L leucine, and 0.007 g/L para-aminobenzoic acid. Unless otherwise indicated, rapamycin, concanamycin A, and indole-3-acetic acid (auxin) were added to cultures at final concentrations of 200 nM, 500 nM, and 1 mM, respectively. For serial-dilution growth assays, exponentially growing yeast cells were all set to a density of 1 OD_600_ / mL in water and five ten-fold serial dilutions were created. 3 μl of each dilution was spotted onto the indicated nutritional media plus agar plates. Unless otherwise indicated, the growth assays were performed at 30°C and images were taken 2 or 3 days later.

### Plasmids

The majority of plasmids generated for this study were created through a similar process that involved insertion of PCR-amplified genes of interest (GOI) into the yeast expression pRS series plasmids (Sikorski and Hieter, 1989) via Gibson assembly (Gibson et al., 2009) and expression was driven from the inclusion of the GOI endogenous promoter or a low expression promoter (*CPY* promoter for N-terminal GFP constructs or *COX8* promoter for MTS-containing constructs). All oligo pairs used in the amplification of GOIs are listed in Table S2 and all plasmids used in this study are listed in Table S3. All plasmids were checked for sequence errors through the ORFs of GOIs, checked for protein expression, and often the function of expressed proteins were analyzed. A brief description for the construction of all *OAC1* containing plasmids is provided for detail. The *OAC1* ORF plus ∼500 base pairs (bp) upstream (5’UTR) and downstream (3’UTR) were PCR-amplified from BY4741 genomic DNA and contained additional 20-25bp flanking regions that were homologous to the DNA regions directly flanking the EcoR1 recognition site found in the MCS of pRS416 and pRS426 plasmids. This *OAC1* amplicon was inserted into both pRS416 and pRS426 plasmids linearized by EcoR1 digestion and the plasmids were reassembled containing the *OAC1* locus via Gibson assembly. Similarly, *OAC1* was fused to GFP in frame at the N-terminus by inserting via Gibson assembly the *OAC1* ORF plus 500bp of its 3’UTR into pGO36 and pGO35, both of which were linearized by digestion with EcoR1. Both pGO36 and pGO35 express GFP from the *CPY* promoter in pRS416 and pRS426, respectively, and were gifts from Greg Odorizzi. The *OAC1-yeGFP ORF* was PCR amplified from AHY8509 genomic DNA and contained ∼500bp upstream (5’UTR) and the *ADH1* terminator sequence and was also inserted into the EcoR1 cut-site of pRS416 and pRS426 plasmids. Alternatively, only the *OAC1-yeGFP ORF* (minus the first start codon) and the *ADH1* terminator sequence were PCR-amplified from AHY8509 genomic DNA and fused behind a PCR-amplicon that contained ∼500bp of the *COX8* 5’UTR and the Cox8 MTS (codons 1-27), both of which were inserted into the EcoR1 cut-site of pRS416 by tandem Gibson assembly. Plasmids for GPD-driven overexpression of OMM proteins, *OAC1*, *AAC1*, or GFP-TA proteins were generated by gateway mediated transfer of the corresponding ORF (Harvard Institute of Proteomics) from pDONR201/221 into pAG306GPD-ccdB chromosome I or pAG413GFP-ccdb (Hughes and Gottschling, 2012) using Gateway LR Clonase II Enzyme mix (ThermoFisher) according to the manufacturer’s instructions. To integrate the resulting expression plasmid into yeast chromosome I (199456-199457), pAG306GPD-ORF chromosome I plasmids were digested with NotI.

### MDC Assays

For MDC assays, overnight log-phase cell cultures were grown in the presence of dimethyl sulfoxide (DMSO) or the indicated drug for two hours. For MDC assays with cells containing plasmids, overnight log-phase yeast cultures grown in selective SD medium were back-diluted to an OD_600_=0.1-0.2 in YPAD medium and allowed to grow for at least 4 hours prior to MDC induction. Prior to visualization, cells were harvested by centrifugation, washed once, and resuspended in 100mM HEPES containing 5% glucose. Subsequently, yeast were directly plated onto a slide at small volumes to allow the formation of a monolayer and optical z-sections of live yeast cells were acquired with a ZEISS Axio Imager M2 or for super-resolution confocal fluorescence microscopy images a ZEISS LSM800 with Airyscan was used. The percent cells with MDCs were quantified in each experiment at the two-hour time point. All quantifications show the mean ± standard error from three biological replicates with *n* = 100 cells per experiment. MDCs were identified as Tom70-positive, Tim50-negative structures that were enriched for Tom70 versus the mitochondrial tubule. In MDC colocalization assays, MDCs were identified as large, Tom70-enriched, spherical structures prior to assessing the co-localization of different proteins of interest.

### Fluorescence Microscopy

Fluorescence microscopy was performed as described in English et al., 2020. In brief, optical z-sections of live yeast cells were acquired with a ZEISS Axio Imager M2 equipped with a ZEISS Axiocam 506 monochromatic camera, 100x oil-immersion objective (plan apochromat, NA 1.4) or 63x oil-immersion objective (plan apochromat, NA 1.4) or a ZEISS LSM800 equipped with an Airyscan detector, 63x oil-immersion objective (plan apochromat, NA 1.4). Crudely purified mitochondria were immobilized on glass slides coated with poly-L-lysine and optical z-sections were acquired with a ZEISS LSM800 equipped with an Airyscan detector, 63x oil-immersion objective (plan apochromat, NA 1.4). Widefield images were acquired with ZEN (Carl Zeiss) and processed with Fiji (Schindelin et al., 2012). Super-resolution images were acquired with ZEN (Carl Zeiss) and processed using the automated Airyscan processing algorithm in ZEN (Carl Zeiss) and Fiji. Individual channels of all images were minimally adjusted in Fiji to match the fluorescence intensities between channels for better visualization. Line-scan analysis was performed on non-adjusted, single z-sections in Fiji.

### Protein Preparation and Immunoblotting

For western blot analysis of protein levels, yeast cultures were grown to log-phase (OD_600_= 0.5-1) and 2 OD_600_ cell equivalents were isolated by centrifugation, washed with dH_2_O and incubated in 0.1 M NaOH for five minutes at RT. Subsequently, cells were reisolated by centrifugation at 16,000 × g for ten minutes at 4°C and lysed for five minutes at 95°C in lysis buffer (10 mM Tris pH 6.8, 100 mM NaCl, 1 mM EDTA, 1 mM EGTA, 1% SDS and containing cOMPLETE protease inhibitor cocktail (Millipore Sigma)). Upon lysis, samples were denatured in Laemmli buffer (63 mM Tris pH 6.8, 2% SDS, 10% glycerol, 1 mg/ml bromophenol blue, 1% β-mercaptoethanol) for five minutes at 95°C. To separate proteins based on molecular weight, equal amounts of protein were subjected to SDS polyacrylamide gel electrophoresis and transferred to nitrocellulose membrane (Millipore Sigma) by semi-dry transfer. Nonspecific antibody binding was blocked by incubation with Tris buffered saline + 0.05% Tween-20 (TBST) containing 10% dry milk (Sigma Aldrich) for one hour at RT. After incubation with the primary antibodies at 4°C overnight, membranes were washed four times with TBST and incubated with secondary antibody (goat-anti-rabbit or donkey-anti-mouse HRP-conjugated,1:5000 in TBST + 10% dry milk, Sigma Aldrich) for 1 hour at RT. Subsequently membranes were washed twice with TBST and twice with TBS, enhanced chemiluminescence solution (Thermo Fisher) was applied and the antibody signal was detected with a BioRad Chemidoc MP system. All blots were exported as TIFFs and cropped in Adobe Photoshop CC.

### Protein Localization within Swollen Mitochondria

All yeast strains examined for protein localization within swollen mitochondria contained Mdm12 C-terminally fused to AID-6xFLAG from the constructs described in Morawska and Ulrich, 2013. Subsequently, GPD1-OsTIR1 was integrated into the *LEU2* locus, using the plasmid pNH605-pGPD1-osTIR1 digested with Swa1 as described in Chan et al., 2018. Auxin-induced protein degradation was performed essentially as described in John Peter et al., 2022, except 3-indole acetic acid (auxin) was added to a final concentration of 1 mM. After a 3-hour incubation, cells were prepped for fluorescence microscopy analysis and the auxin-induced degradation of Mdm12 and swelling of mitochondria was visually confirmed as shown in John Peters et al., 2022.

### Isolation of Yeast Mitochondria

Crudely purified mitochondria were isolated from yeast cells as described in Schuler et al., 2021. Briefly, yeast were grown overnight in log-phase to an OD_600_= 0.5-1 as described above, then isolated by centrifugation, washed with dH_2_O and the pellet weight was determined. Cells were then resuspended in 2 ml/g pellet dithiothreitol (DTT) buffer (0.1 M Tris, 10 mM DTT, pH 9.4) and incubated for 20 minutes at 30°C under constant shaking. After re-isolation by centrifugation, DTT treated cells were washed once with zymolyase buffer (1.2 M sorbitol, 20 mM K_2_HPO_4_, pH 7.4 with HCl) and cell walls were digested for 30 minutes at 30°C under constant shaking in 7 ml zymolyase buffer / g cell pellet containing 1 mg zymolyase 100T / g cell pellet. After zymolyase digestion, cells were reisolated by centrifugation, washed with zymolyase buffer and lysed by mechanical disruption in 6.5 ml/g pellet homogenization buffer (0.6 M sorbitol, 10 mM Tris pH 7.4, 1 mM ethylenediaminetetraacetate (EDTA) pH 8.0 with KOH, 0.2% BSA, 1 mM phenylmethylsulfonylfluoride) at 4°C. Cell debris were removed from the homogenate twice by centrifugation at 5000 × g for five min at 4°C and mitochondria were pelleted at 14000 × g for 15 min at 4°C. The mitochondrial pellet was resuspended in SEM buffer (250 mM sucrose, 1 mM EDTA pH 8.0 with KOH, 10 mM 3-(*N*-morpholino)- propansulfonic acid pH 7.2), reisolated by differential centrifugation as described above, resuspended in SEM buffer and mitochondria were shock frozen in liquid nitrogen and stored at −80°C.

### Protease Protection and Carbonate Extraction

For protease protection assays, 50 µg of crudely purified mitochondria were incubated on ice with or without proteinase K (PK) under different conditions: mitochondria were stabilized in an iso-osmolar buffer (SEM buffer), the OMM was ruptured by hypo-osmolar swelling (EM buffer: 1 mM EDTA, 10 mM MOPS/KOH, pH 7.2), or mitochondria were lysed by addition of 1% Triton X-100 (TX) to SEM buffer. Subsequently, PK activity was inhibited by the addition of 2 mM PMSF, mitochondria were pelleted by centrifuging at 20,000 x *g* at 4°C, and analyzed by SDS-PAGE and immunoblotting as described above. For carbonate extraction experiments, 100 µg of mitochondria were incubated on ice with 0.1 M Na_2_CO_3,_ pH 11 for 30 min. Subsequently, mitochondrial membranes were pelleted by centrifuging at 126,000 x *g* for 45 min at 4°C, and analyzed by SDS-PAGE and immunoblotting as described above.

## QUANTIFICATION AND STATISTICAL ANALYSIS

The number of replicates, what *n* represents, and dispersion and precision measures are indicated in the Figure Legends. In general, quantifications show the mean ± standard error from three biological replicates with *n* = 100 cells per experiment. In experiments with data depicted from a single biological replicate, the experiment was repeated with the same results.

## ACKNOWLEDGEMENTS

We thank members of the A.L. Hughes group for discussion and manuscript comments. We thank members of Janet. M. Shaw laboratory for providing reagents, mitochondrial antibodies, and support early on in the project. We would like to also thank Greg Odorizzi for plasmid gifts and Maya Schuldiner for gifting the yeast SWAp-Tag library described in Weill et al., 2018. Zachary Wilson, PhD, was supported by an American Heart Association Postdoctoral Fellowship, 20POST35200110, and supported by the American Cancer Society–Give Mas, Live Mas Southern Multifoods Postdoctoral Fellowship, PF-20-018- 01-CCG. Research was further supported by National Institutes of Health grant GM119694 to A.L.H.

## AUTHOR CONTRIBUTIONS

Conceptualization, Z.N.W. and A.L.H.; methodology, Z.N.W, S.B., and M.W.; formal analysis, Z.N.W.; investigation, Z.N.W. S.B., and M.W.; writing – original draft, Z.N.W; writing – review and editing, Z.N.W. and A.L.H.; visualization, Z.N.W.; supervision, A.L.H.; funding acquisition, Z.N.W. and A.L.H.

## DECLARATION OF INTERESTS

The authors declare no competing interests.

## CONTACT FOR REAGENT AND RESOURCE SHARING

Further information and requests for resources and reagents should be directed to and will be fulfilled by the Lead Contact, Adam Hughes. All unique/stable reagents generated in this study are available from the Lead Contact without restrictions.

## SUPPLEMENTAL FIGURE LEGENDS

**Figure S1.**
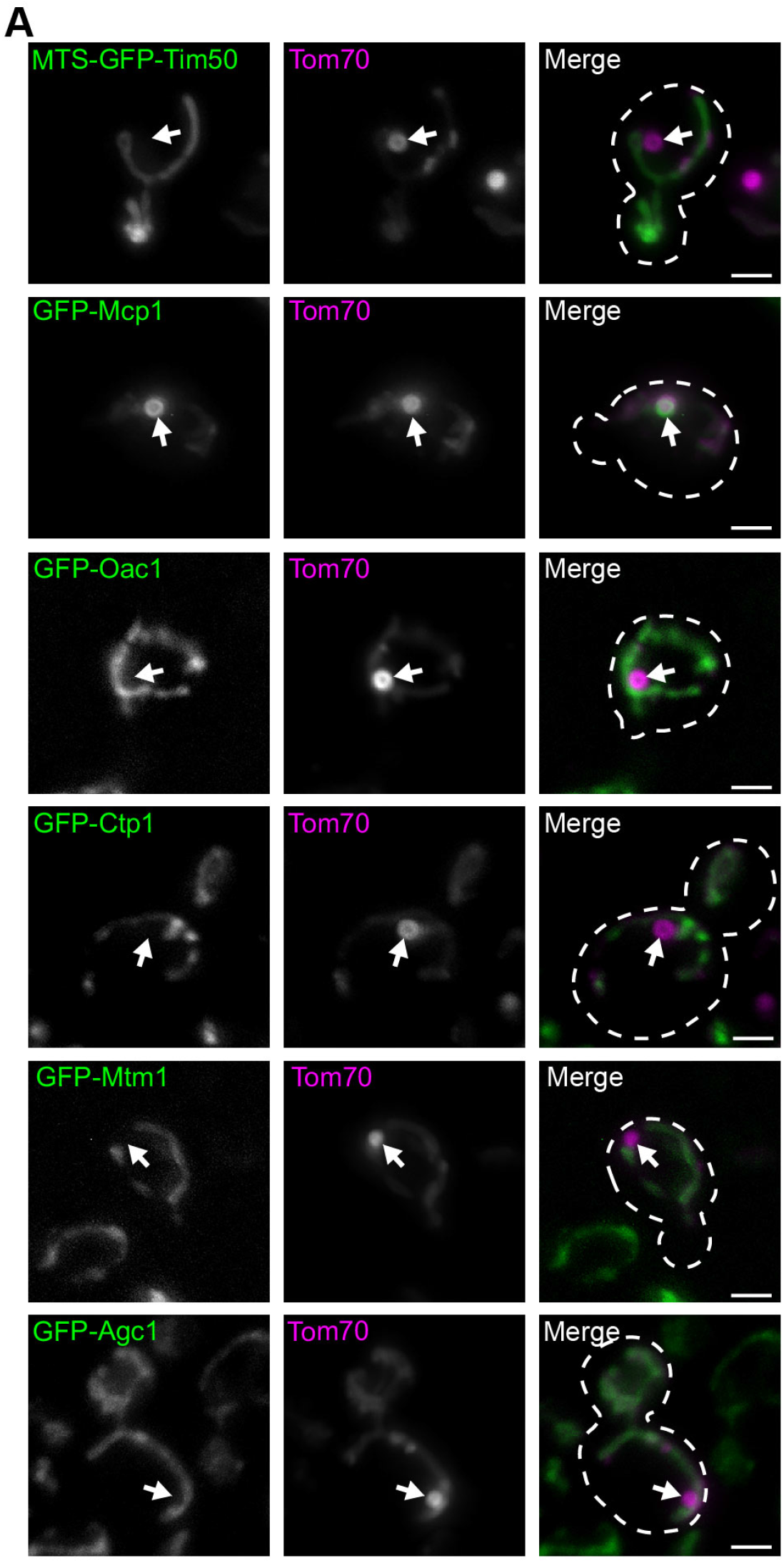
N-terminally tagged mitochondrial metabolite carriers are excluded from MDCs. (A) Representative max projections of widefield fluorescence microscopy images of yeast expressing Tom70-mCherry and the indicated sfGFP-tagged strains examined from the N-terminal SWAp-Tag (N-SWAT) library all treated with 200 nM rapamycin. MDCs are indicated by the white arrows. Scale bar= 2 µm.

**Figure S2.**
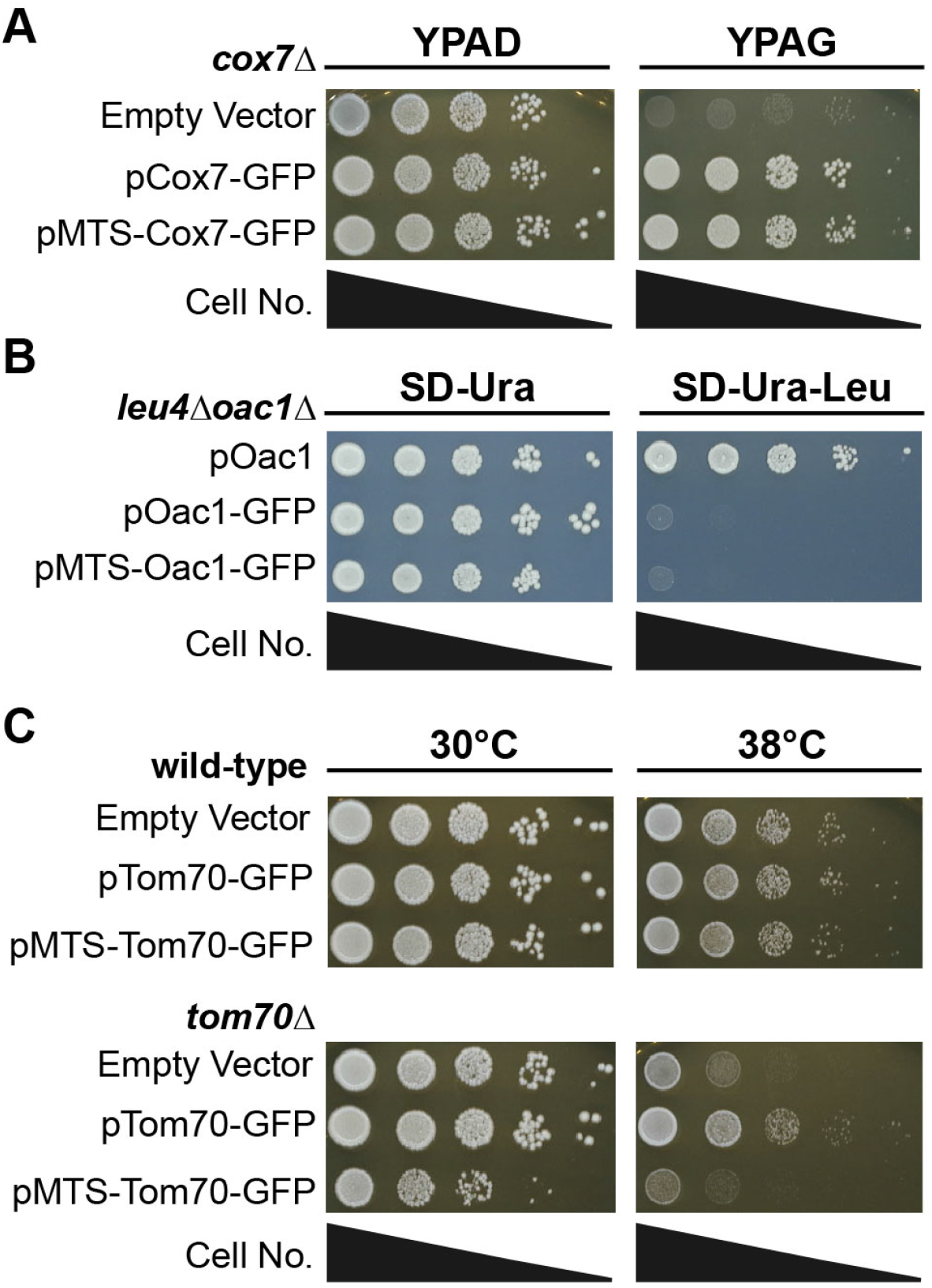
Functional analysis of MTS-containing fusion proteins. (A) Cell growth spot assays of *cox7*Δ yeast transformed with the indicated Cox7 constructs or empty vector expressed from a low-copy (pRS416) plasmid serially-diluted and plated on either rich media agar plates containing dextrose (YPAD) or glycerol (YPAG). (B) Cell growth spot assays of *leu4*Δ*oac1*Δ yeast transformed with the indicated Oac1 constructs expressed from a low-copy (pRS416) plasmid serially-diluted and plated on SD agar plates lacking either uracil (SD-Ura) or uracil and leucine (SD-Ura-Leu). (C) Cell growth spot assays of wild-type or *tom70*Δ yeast transformed with the indicated Tom70 constructs or empty vector expressed from a low-copy (pRS416) plasmid serially-diluted and plated on YPAD media agar plates and grown at either 30°C or 38°C.

**Figure S3.**
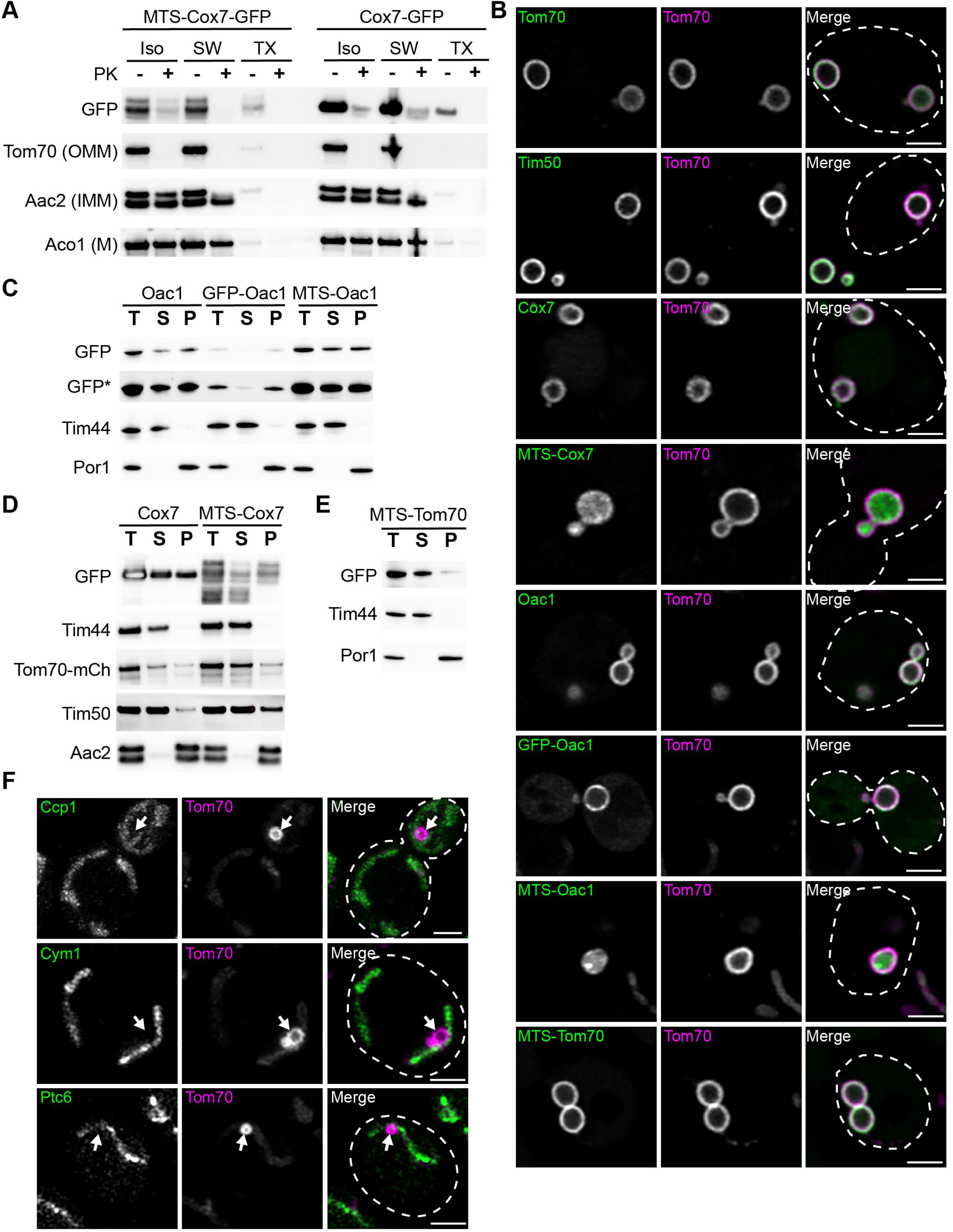
MDC cargo are localized to the outer mitochondrial membrane. (A) Immunoblot of a protease protection assay on isolated mitochondria from yeast cells expressing Tom70-mCherry and either Cox7-GFP or MTS-Cox7-GFP. The extent of proteinase K (PK) protection was analyzed through immunoblot with antibodies specific to either GFP, mCherry for Tom70-mCherry (OMM), Aac2 (IMM), or Aco1 (matrix, M). ISO= iso-osmolar buffer, SW= hypo-osmolar swelling, TX= 1% Triton X-100. (B) Super-resolution confocal fluorescence microscopy images of swollen mitochondria in yeast cells expressing Tom70-mCherry, Mdm12-6xFLAG-AID and OsTir1 treated with 1 mM auxin for 3 hours. These yeast cells were also transformed with a low-copy plasmid (pRS416) expressing the indicated GFP-tagged proteins. Scale Bar= 2 µm. (C-E) Mitochondrial membrane integration was analyzed through a carbonate extraction assay. Mitochondria isolated from yeast cells expressing either Cox7-GFP (C), MTS-Cox7-GFP (C), Oac1-GFP (D), GFP-Oac1 (D), MTS-Oac1-GFP (D), or MTS-Tom70-GFP (E) were treated with 100 mM Na_2_CO_3_ pH 11 to transform mitochondrial membranes into membrane sheets. Membranes were precipitated through ultracentrifugation and the extent that proteins remained in the membrane pellet fraction (P) compared to the supernatant (S) was determined through immunoblot with antibodies specific to either GFP, mCherry for Tom70-mCherry (OMM protein), Tim50 (IMM protein), Aac2 (polytopic IMM protein), Por1 (OMM protein) or Tim44 (membrane-associated matrix protein). A total (T) untreated sample of isolated mitochondria were also analyzed as a control for input. * indicates a longer exposure. (F) Super-resolution confocal fluorescence microscopy images of rapamycin-induced MDC formation in yeast expressing Tom70-mCherry and the indicated intermembrane space proteins C-terminally tagged to GFP. MDCs are indicated by the white arrows. Scale bar= 2 µm. Images from Ccp1-GFP cells are max projections, the other images are from a single z-slice.

**Figure S4.**
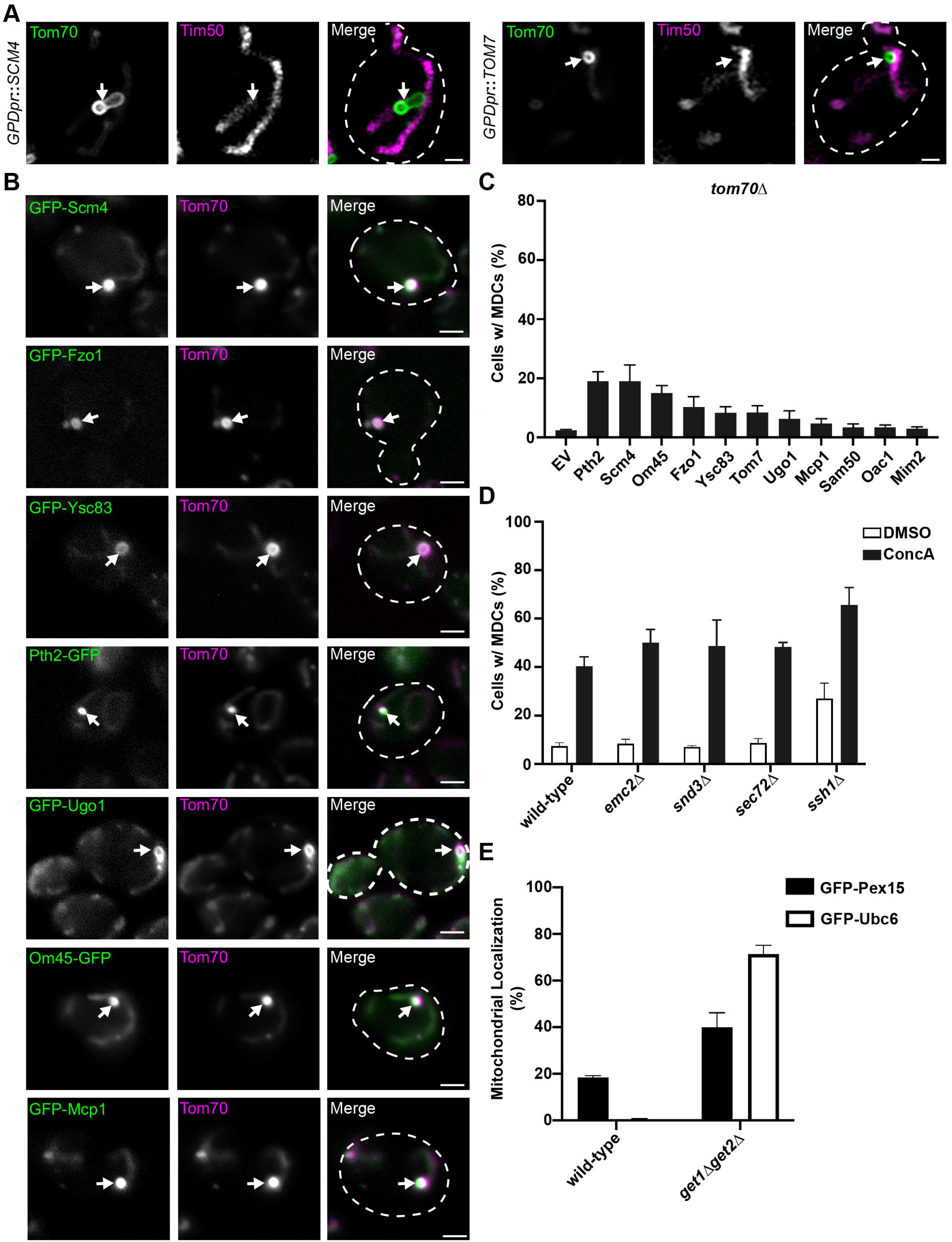
Overloading the OMM with membrane proteins induces MDC formation. (A) Super-resolution confocal fluorescence microscopy images of cells expressing Tom70-GFP and Tim50-mCherry and constitutively overexpressing Scm4 or Tom7. MDCs are indicated by the white arrows. Scale Bar= 1 µm. (B) Representative max projections of widefield fluorescence microscopy images of MDC formation of yeast expressing Tom70-mCherry and the indicated OMM proteins tagged with GFP. MDCs are indicated by the white arrows. Scale bar= 2 µm. (C) Quantification of the percent of cells forming MDCs in *tom70*Δ cells constitutively overexpressing the indicated proteins. Error bars show mean ± standard error of three replicates, *n* ≥ 100 cells per replicate. (D) Quantification of the percent of cells forming MDCs in the indicated yeast strains treated with DMSO (vehicle control) or 500nM ConcA. Error bars show mean ± standard error of three replicates, *n* ≥ 100 cells per replicate. (E) Quantification of the percent of cells containing mitochondrial localized GFP-Pex15 or GFP-Ubc6 in the indicated yeast strains. Error bars show mean ± standard error of three replicates, *n* ≥ 100 cells per replicate.

## Notes

### Competing Interest Statement

The authors have declared no competing interest.

