## Supplementary material for "Mitochondrial-Derived Compartments Remove Surplus Proteins from the Outer Mitochondrial Membrane": Table S3

**Table S2. Bacterial strains, chemicals, plasmids, and software used in this study**

| **Reagent or resource** | | **Source** | | **Identifier** |
| --- | --- | --- | --- | --- |
| **Antibodies** | | | | |
| Mouse monoclonal anti-GFP clones 7.1 and 13.1 (dilution 1:1000) | Roche | | Cat # 11814460001; RRID:AB_390913 | |
| Mouse monoclonal anti-PGK1 clone 22C5D8 (dilution 1:5000) | Abcam | | Cat # ab113687; RRID:AB_10861977 | |
| Mouse monoclonal anti-Por1 clone 16G9E6BC4 (dilution 1:1000) | Abcam | | RRID:AB_10865182; Cat # ab110326 | |
| Rabbit polyclonal anti-Aco1 (dilution 1:2000) | Shaw Lab | | N/A | |
| Rabbit polyclonal anti-Aac2 (dilution 1:5000) | Shaw Lab | | N/A | |
| Rabbit polyclonal anti-Tim44 (dilution 1:1000) | Pfanner Lab | | N/A | |
| Rabbit polyclonal anti-Tim50 (dilution 1:1000) | Pfanner Lab | | N/A | |
| **Bacterial strains** | | | | |
| *Escherichia coli* DH5α | | N/A | | N/A |
| *S. cerevisiae* ORF collection (pDONR201/221) | | Harvard Institute of Proteomics | | N/A |
| One Shot™ *ccd*B Survival™ 2 T1^R^ Competent Cells | | Thermo Fisher | | Cat#A10460 |
| **Chemicals, peptides, and recombinant proteins** | | | | |
| 3-Indoleacetic acid (Auxin) | | Sigma-Aldrich | | Cat # I3750 ; CAS # 87-51-4‎ |
| Casamino acids | | US Biological | | Cat # 0012501A; CAS # 65072-00-6 |
| Concanamycin A | | Santa Cruz Biotechnology | | Cat # sc-202111; CAS # 80890-47-7 |
| Concanavalin A | | Sigma-Aldrich | | Cat # L7647; CAS # 11028-71-0 |
| Dimethyl sulfoxide (DMSO) | | Sigma-Aldrich | | Cat # D2650; CAS # 67-68-5 |
| DTT | | Gold Biotechnology | | Cat # DTT10; CAS # 27565-41-9 / 3483-12-3 |
| Lyticase | | Sigma-Aldrich | | Cat # L2524; CAS #  37340-57-1 |
| Rapamycin | | LC Laboratories | | Cat # R-5000; CAS # 53123-88-9 |
| Zymolyase 100T | | Amsbio | | Cat # 120493-1; CAS #37340-57-1 |
| Critical commercial assays | | | | |
| Bicinchoninic Acid Protein Assay | | Thermo Fisher | | Cat # 23227 |
| Gateway LR Clonase II Enzyme Mix | | Thermo Fisher | | Cat # 11791020 |
| Gibson Assembly Master Mix | | New England Biolabs | | Cat # E2611 |
| **Plasmids** | | | | |
| Plasmid: pAG306GPD-AAC1 chr 1 | | Schuler et al. 2021 | | B4018 |
| Plasmid: pAG306GPD-ccdB chr 1 | | Hughes and Gottschling, 2012 | | B3681 |
| Plasmid: pAG306GPD-ECM19 chr 1 | | This Study | | B4019 |
| Plasmid: pAG306GPD-FIS1 chr 1 | | This Study | | pAH464 |
| Plasmid: pAG306GPD-FZO1 chr 1 | | This Study | | B4020 |
| Plasmid: pAG306GPD-MCP1 chr 1 | | English et al. 2020 | | B4021 |
| Plasmid: pAG306GPD-MIM2 chr 1 | | This Study | | B4022 |
| Plasmid: pAG306GPD-OAC1 chr 1 | | Schuler et al. 2021 | | B4023 |
| Plasmid: pAG306GPD-OM45 chr 1 | | This Study | | B4024 |
| Plasmid: pAG306GPD-POR1 chr 1 | | This Study | | B3685 |
| Plasmid: pAG306GPD-PTH2 chr 1 | | This Study | | B4025 |
| Plasmid: pAG306GPD-SAM50 chr 1 | | This Study | | B4026 |
| Plasmid: pAG306GPD-SCM4 chr 1 | | This Study | | B4027 |
| Plasmid: pAG306GPD-SEN54 chr 1 | | This Study | | B4028 |
| Plasmid: pAG306GPD-TMH11 chr 1 | | This Study | | B4032 |
| Plasmid: pAG306GPD-TOM20 chr 1 | | Schuler et al. 2021 | | B3773 |
| Plasmid: pAG306GPD-TOM6 chr 1 | | This Study | | B4029 |
| Plasmid: pAG306GPD-TOM7 chr 1 | | This Study | | B4030 |
| Plasmid: pAG306GPD-TOM70 chr 1 | | Schuler et al. 2021 | | N/A |
| Plasmid: pAG306GPD-TOM71 chr 1 | | This Study | | B4031 |
| Plasmid: pAG306GPD-UGO1 chr 1 | | This Study | | B3774 |
| Plasmid: pAG306GPD-YSC83 chr 1 | | This Study | | B4033 |
| Plasmid: pAG413-GPD-EGFP-PEX15 | | This Study | | pAH425 |
| Plasmid: pAG413-GPD-EGFP-UBC6 | | This Study | | pAH428 |
| Plasmid: pHLUM | | Mülleder *et al*., 2012; Addgene | | Plasmid # 40276 |
| Plasmid: pHYG-AID*-6FLAG | | Morawska and Ulrich, 2013 Addgene | | Plasmid # 99519 |
| Plasmid: pKT127-mCherry | | Daniel Gottschling (Calico) | | N/A |
| Plasmid: pKT128 | | Sheff and Thorn, 2004; Addgene | | Plasmid # 8729 |
| Plasmid: pRS305 | | Sikorski and Hieter, 1989 | | N/A |
| Plasmid: pRS306 | | Sikorski and Hieter, 1989 | | N/A |
| Plasmid: pRS400 | | Daniel Gottschling (Calico) | | N/A |
| Plasmid: pRS40Hyg | | Daniel Gottschling (Calico) | | N/A |
| Plasmid: pRS40Nat | | Daniel Gottschling (Calico) | | N/A |
| Plasmid: pRS416-COX7-yEGFP | | This Study | | B3746 |
| Plasmid: pRS413-COX7-TIM50-yEGFP | | This Study | | B3815 |
| Plasmid: pRS413-COX8-TIM50-yEGFP | | This Study | | B3830 |
| Plasmid: pRS416-GFP-OAC1 | | This Study | | B3752 |
| Plasmid: pRS426-GFP-OAC1 | | This Study | | B3753 |
| Plasmid: pRS416-MTS-COX7-yEGFP | | This Study | | B3748 |
| Plasmid: pRS413-MTS-COX7-TIM50-yEGFP | | This Study | | B3762 |
| Plasmid: pRS416-MTS-OAC1-yEGFP | | This Study | | B3766 |
| Plasmid: pRS416-MTS-TOM70-yEGFP | | This Study | | B3767 |
| Plasmid: pRS416-OAC1 | | This Study | | B3822 |
| Plasmid: pRS426-OAC1 | | This Study | | B3823 |
| Plasmid: pRS416-OAC1-yEGFP | | This Study | | B3990 |
| Plasmid: pRS426-OAC1-yEGFP | | This Study | | B3992 |
| Plasmid: pRS413-TIM50-yEGFP | | This Study | | B3748 |
| Plasmid: pRS416-TIM50 | | This Study | | B3831 |
| Plasmid: pRS416-TOM70-yEGFP | | This Study | | B3750 |
| **Software and algorithms** | | | | |
| FIJI | | Schindelin *et al*., 2012 | | Version 1 |
| Prism | | GraphPad Software, Inc. | | Version 9 |
| SnapGene | | GSL Biotech | | Version 4.2 |
| ZEN Blue Edition | | Carl Zeiss Microscopy | | Version 2.6 |
